# PATO: Pangenome Analysis Toolkit

**DOI:** 10.1101/2021.01.30.428878

**Authors:** Miguel D. Fernández-de-Bobadilla, Alba Talavera-Rodríguez, Lucía Chacón, Fernando Baquero, Teresa M. Coque, Val F. Lanza

**Affiliations:** Department of Microbiology, University Hospital Ramón y Cajal, IRYCIS, Madrid, Spain; Department of Infectious Diseases, University Hospital Ramon y Cajal, IRYCIS, Madrid, Spain; Bioinformatics Unit, University Hospital Ramón y Cajal, IRYCIS, Madrid, Spain

## Abstract

**Motivation:** Comparative genomics is a growing field but one that will be eventually overtaken by sample size studies and the increase of available genomes in public databases. We present the Pangenome Analysis Toolkit (PATO) designed to simultaneously analyze thousands of genomes using a desktop computer. The tool performs common tasks of pangenome analysis such as core-genome definition and accessory genome properties and includes new features that help characterize population structure, annotate pathogenic features and create gene sharedness networks. PATO has been developed in R to integrate with the large set of tools available for genetic, phylogenetic and statistical analysis in this environment.

**Results:** PATO can perform the most demanding bioinformatic analyses in minutes with an accuracy comparable to state-of-the-art software but 20–30x times faster. PATO also integrates all the necessary functions for the complete analysis of the most common objectives in microbiology studies. Lastly, PATO includes the necessary tools for visualizing the results and can be integrated with other analytical packages available in R.

**Availability:** The source code for PATO is freely available at https://github.com/irycisBioinfo/PATO under the GPLv3 license.

**Supplementary information:** Supplementary data are available at Bioinformatics online

## 1 Introduction

Comparative genomics and microbial genomics are two of the fastest-growing research fields of the last decade. The unstoppable reduction in sequencing costs has made the sequencing of large numbers of strains affordable. As a consequence, there is an increasing need for methods of analyzing not just hundreds but thousands of sequences using conventional computer resources. There are numerous solutions for analyzing pangenomes (the entire set of genes/genomes), such as Roary (Page *et al.*, 2015), OrthoMCL (Li *et al.*, 2003), PIRATE (Bayliss *et al.*, 2019), PanX (Ding *et al.*, 2018) and Panaroo (Tonkin-Hill *et al.*, 2020). We present the Pangenome Analysis Toolkit (PATO), an R package that helps manage and analyze large data sets. PATO not only can determine the pangenome but also contains numerous options for analyzing the pangenome, the core genome and the accessory genome and visualizing the results. The significant differences between rival methods makes it difficult to perform reliable comparisons. PATO includes tools for most of the common tasks in comparative genomics: phylogenetics, population structure, annotation and representation. PATO handles the main file formats and is freely available for Unix and MacOS systems.

## 2 Implementation

We developed PATO as an R package for several reasons. First, R is currently one of the main environments for large data analysis. Second, R (along with the Bioconductor repository) has a large collection of packages related to genomics, population analysis, data visualization and, of course, statistical analysis. Third, R is free, multiplatform and allows for parallel computing. PATO contains internal R functions but also uses external software. The PATO core functions use MMSeqs2 (Steinegger and Söding, 2018), MASH (Ondov *et al.*, 2016) and Minimap2 (Li, 2018). To avoid incompatibility issues, PATO includes the executables of these software applications (Unix and MacOS). We have implemented multithreading properties as much as possible in the PATO functions. In addition to these executables, PATO contains a number of auxiliary programs such as *bedtools* (Quinlan, 2014), *k8* (https://github.com/attractivechaos/k8) and a number of customized Perl scripts.

PATO draws from other databases for annotating sequences associated with pathogenicity (Virulence Factor DataBase) (Chen *et al.*, 2016) and resistance to antibiotics (ResFinder) (Bortolaia *et al.*, 2020). PATO also includes a customized database of reference and representative bacterial genomes from RefSeq (all of which were last updated in October 2020).

PATO is built with a set of objects (S3 objects) and functions to increase the robustness and compatibility among the functions (See *Functions and Classes* in the Supplementary Material). The output of a function can therefore be the input of another function. Some functions use temporal files and folders, which PATO can re-use when necessary to save computational time.

For common processes such as determining the core genome and the core-snp-genome, PATO has proven to be 35 times and 23 times faster than Roary and Snippy, respectively, which currently represent the state-of-the-art software for this purpose (see *Benchmarking* section in Supplementary Data).

## 3 Features

PATO can handle most of the sequence files formats, such as whole-genome nucleotide (fna), nucleotide features or CDS (ffn) and proteome files (faa). In addition, PATO includes a function to load GFF files, provided they contain the FASTA sequence at the end of the file. This format is one of the outputs of PROKKA (Seemann, 2014), which is widely employed in bacterial annotation.

We have divided the PATO functions into four sections: Quality Control, Main Functions, Analysis and Visualization.

Quality Control implements the functions to filter out outliers and identify species. It is sometimes possible to find sequences that should not belong in the sample, both in public databases and in our own datasets. PATO can find outliers among the dataset and classify each with the most related species. We have included a customized reference database with all the reference and representative bacterial genomes from RefSeq. Another common issue is redundancies in the datasets. Although there is a frequent need for a homogeneous representation of a species, databases often have an over-representation of certain phylogenetic branches (e.g., isolates from outbreaks). PATO implements functions to create a non-redundant representation of genomes.

The main functions create objects that other functions use (analysis, visualization and quality control). Most of the main functions are wrappers for external software. These functions first execute the external software and then load, transform and build the corresponding R object with the results. PATO depends on MMSeqs2 to cluster family proteins and/or genes to build the core-genome and the accessory genome. MMSeqs2 is one of the fastest and most accurate methods for clustering genes and proteins. Moreover, the implementation of MASH helps quickly estimate the similarity among the sequences. Lastly, PATO employs Minimap2 to create a whole-genome alignment and build the core snp genome (all common polymorphic positions of the set of genomes). In addition to Minimap2, PATO employs the software application *paftools.js* to perform the variant calling of the Minimap2 results following the pipeline proposed by the Minimap2 developers. PATO has a set of functions to create and analyze a set of pangenomes. The idea is to treat each pangenome as an individual sample and then analyze all of them as a whole. Depending on the dataset size, PATO implements two pipelines, one for datasets smaller than 5000 genomes and the other for datasets over 5000 genomes. We have created a set of functions that build groups of genomes and convert them into pangenomes or use pangenomes that have been pre-defined by the user. Once the pangenomes have been obtained, we create an *accnet* object and analyze it with common tools. PATO makes it possible to analyze tens of thousands of genomes in a single study (see Supplementary “Pangenome Analysis”)

PATO also implements the latest version of AccNET (Lanza *et al.*, 2017) and all the necessary functions to analyze it. However, to visualize this kind of data, we recommend using Gephi (Bastian *et al.*, 2009), which helps create networks with hundreds of genomes (see Fig. 8 of Supplementary Material).

One of the reasons to use R is the broad availability of packages for data visualization. PATO implements several functions to visualize the results. We have included common features such as core, accessory and pangenome distribution, beta diversity (multidimensional scaling through UMAP (Becht *et al.*, 2018)) and similarity trees (MASH tree or accessory tree). However, we have developed a novel way to represent the population structure. Using a combination of k-nearest neighbor and spanning tree concepts, we have created the K-Nearest Neighbor

Network visualization. This function selects the best K neighbors for each genome and creates a network. Our novel approach includes the option of avoiding reciprocal connections, which means that each genome will have a set of the nearest neighbors that are not yet connected to itself (see *Population Structure* section in Supplementary Material).

Within the analysis type functions, we have functions for the statistical analysis of the accessory genome and gene association (coincidences, twins, singles), clustering functions (both for the whole genome and for the accessory genome), plasmid sequence extraction, SNP distribution analysis and feature annotation (antibiotic resistance genes and virulence factors).

We have implemented a previous pipeline to analyze enrichment genes among accessory genomes (Fernández-de-Bobadilla *et al.*, 2020). This function determines which genes are over-represented in a group compared with the population (i.e., the total set of genomes analyzed). PATO also implements functions to build an SNP matrix. We have also included a function that uses *mlplasmids* (Arredondo-Alonso *et a.*, 2018) packages (which need to be installed) to extract the plasmidome of the data set. Lastly, PATO has a set of functions to export data to other external software applications for network analysis, visualization and phylogenetic reconstruction.

## 4 Conclusion

PATO is a package that can perform not only the most routine and common tasks in comparative genomics but also advanced and novel pangenome analysis. To this end, we have employed the most efficient tools currently available and integrated them into a Big Data analysis environment. PATO can analyze thousands of genomes in a few hours and integrate common tools used in phylogenetic and statistical analysis.

In the Supplementary Material, we show how PATO can perform an analysis of outbreak, characterize the population structure of a microbial species and perform gene flow analysis on genome datasets corresponding to multistrain and multispecies collections.

