## Supplementary Material for "PATO: Pangenome Analysis Toolkit"

2021-01-29

### Introduction

PATO is an R package designed to analyze pangenomes (set of genomes) from the same or different species (intra/inter species). PATO allows the analysis of population structure, phylogenetics and horizontal gene transfer using in each case the core-genome (set of genes common to all genomes), the accessory genome (set of "dispensable" genes) or the whole genome. PATO uses external software in its core software: MASH, MMSeq2, Minimap2.

This software can handle thousands of genomes using conventional computers without the need of using HPC facilities. We have designed PATO to work with the most common files such as GFF (General Feature Format), FNA (FASTA Nucleotide ), FFN (FASTA Feature Nucleotide) or FAA (FASTA Aminoacid). PATO is able to perform common genomic analysis of phylogenetic, annotation or population structure. Moreover, PATO implements functions of quality control, dataset manipulation and visualization.

### Installation

#### Linux/Unix

PATO uses external binaries so it cannot be included in the CRAN repository. To install PATO, use the `devtools` package.

Sometimes, some dependencies require system packages as *libcurl* *libssl* or *libxml2* (this example is for Ubuntu/Debian based systems):

```
sudo apt install libcurl4-openssl-dev libssl-dev
 >libxml2-dev libmagick++-dev libv8-dev
```

To install `devtool` type:

```
install.packages("devtools")
```

Once you have installed `devtools` you should activate the Bioconductor repository

```r
setRepositories(ind = c(1,2))
## Select options 1 (CRAN) and 2 (Bioconductor Software)
```

Now you can install PATO from GitHub.

```r
devtools::install_github("https://github.com/irycisBioinfo/PATO.git",
                         build_vignettes = TRUE)
```

##### MacOS

In MacOS systems, you should install `devtools` package first.

```r
install.packages("devtools")
```

Once you have installed `devtools` you should activate the Bioconductor repository.

```r
setRepositories(ind = c(1,2))
## Select options 1 (CRAN) and 2 (Bioconductor Software)
```

Now you can install PATO from GitHub.

```r
devtools::install_github("https://github.com/irycisBioinfo/PATO.git",
                         build_vignettes = TRUE)
```

#### Functions and Classes

PATO implements different wrapper functions to use external software. PATO includes `MMSeqs2`, `Mash` and `Minimap2` binaries and other extra software such as `bedtools`, `k8` and perl and javascript scripts.

PATO has been designed as a toolkit. Most of the functions use as input other outputs from functions, creating a complete workflow to analyze large genome datasets. As an R package, PATO can interact with other packages like `ape`, `vegan`, `igraph` or `pheatmap`. We have tried to develop PATO to be compatible with the `tidyverse` packages in order to use `%>%` pipe command and to be compatible with `ggplot2` and extensions (for example, `ggtree`).

##### Classes

In order to create a robust environment, PATO uses specific data classes (S3 objects) to assure the compatibility among functions. Most of the classes are lists of other kinds of objects (data.frame, character vector, matrix…).

PATO has 8 object classes:

- `mash` object
- `mmseq` object
- `accnet` object
- `nr_list` object
- `gff_list` object
- `core_genome` object
- `core_snps_genome` object
- `accnet_enr` object

Other outputs take external objects such as `igraph` or `ggplot`.

The idea to have these objects is to share structured data among functions. These objects can be inspected and some of their data can be used with external function/packages.

##### mash class

A mash class is a list of three elements: table, matrix and path.

The first one, matrix, is a square and symmetric matrix of the distances among genomes. The matrix has the input file names as rownames and colnames

The second one, table is a data.table/data.frame with all the distances as list (long format). The table has the columns c("Source","Target","Dist")

The third one, path is the full pathname to the temporary folder. This temporary folder can be reused in future runs to shorten computing times.

##### mmseq class

A mmseq object is a list of three elements: table and annot and path.

First element, table is a data.table/data.frame with four columns c("Prot_genome", "Prot_Prot", "Genome_genome", "Genome_Prot"). This is the output of MMSeqs2 and describes the clustering of the input genes/proteins. The first column refers to the genome that contain the representative gene/protein of the cluster. The second column is the representative protein of the cluster (i.e. the cluster name). The third one is the genome that includes the gene/protein of the fourth column.

The second element, annot is a data.frame/data.table with the original annotation of all representative gene/protein of each cluster in two columns. The first one Prot_prot is the same as appears in the second column of the table element. The second column is the gene/protein annotation.

The third one, path is the full pathname to the temporary folder. This temporary folder can be reused in future runs to shorten computing times.

##### accnet class

An accnet object is a list of three elements: list, matrix, annot.

First element, list, is a data.frame/data.table with the columns c("Source","Target","degree"). It contains the correspondences among proteins/genes (Target) and genomes (Source) and the degree of the corresponding protein/gene in the entire network. In some cases, for example, when the accnet object comes from the accnet_with_padj function or pangenomes the third column can be different and include information such as the frequency of the protein/gene in the pangenome (in the case of pangenomes), or the p-value of the association between the genome and the protein/gene (in the case of accnet_with_padj)

Second element, matrix, is a presence/absence matrix. It is very important to know that it has a column Source, so in the case of using the matrix out of PATO you should convert this column into rownames (my_matrix <- my_accnet$matrix %>% column_to_rownames("Source"))

Third element, annot is a data.table/data.frame that contains the original annotation of the representative gene/protein of that cluster.

##### nr_list class

nr_list is the output format of the non redundant functions and contains the clustering results of the inputs. A nr_list is a S3 object of three elements: Source,centrality and cluster. nr_list can be coerced to a data.frame of three columns using the as.data.frame function. Element Source has the accession/name of the genomes, centrality has the centrality value of the genome, according to the degrees of vertices, for its cluster. Finally, cluster has the membership of the Source genome.

##### gff_list class

A gff_list is an object that stores the results of load_gff_list() output. It stores the path to the GFF, FNA, FFN and FAA files. A gff_list object can be used as input for classifier, mash, mmseq, `core_snp_genome` and `snps_pairwaise`. It has two elements: `path` and `files`. The `path` contains the pathname to the temporary directory. That directory has four sub-directories (FAA, FNA, FFN and GFF) in which the FASTA files and the GFF are stored.

This temporary folder can be reused in future runs to shorten computing times.

###### `core_genome` and `core_snps_genome`

These are objects produced by the `core_` functions. They contain the alignment information (core or core_snps) and can be exported to FASTA format or used in other functions. `core_genome` has two elements: `core_genome` and `path`. The `core_genome` element is a data.frame with two columns `c("Genome","Seq")` and contains the core-genome alignment. The `path` indicates the temporary directory. `core_snps_genome` has four elements: `alignment`, `bed`, `path` and `reference`. Like `core_genome$core_genome alignment` contains the core-genome alignment. `bed` is the BED table with the conserved regions of the alignment (with the coordinates in the reference base). `reference` indicates the genome chosen to be the `reference`. Finally, `path` is the pathname to the temporary directory.

###### `accnet_enr`

This object is a `data.frame` with the result of `accnet_enrichment_analysis()`. That object can be exported as a `gephi` file with the p-values (adjusted) as a edge-weight. It has the columns:

Target: Protein Source: Genome Cluster: Cluster of genome perClusterFreq: Percentage (%) of the protein in this cluster ClusterFreq: Frequency [0-1] of the protein in this cluster ClusterGenomeSize: Number of genomes in the cluster perTotalFreq: Percentage (%) of the protein in the population TotalFreq: Frequency [0-1] of the protein in the population OddsRatio: Odds ratio of the protein p value: p-value of the hypergeometric test (see `phyper`) padj: Adjusted p-value using `p.adjust` AccnetGenomeSize: Total number of genomes AccnetProteinSize: Total number of proteins Annot: Original functional annotation of the protein.

#### Functions

PATO includes internal functions and other wrap functions that calls external binaries. We split the functions into 4 categories: *Main Functions*, *Analysis Functions*, *Quality Control* and *Visualization*.

##### Main Functions

**`load_gff_list()`** There are several kinds of sequence files: whole genome, CDS or Aminoacid files. PATO can handle all of them (not in all functions). One way to have all the information together is using GFF files. These files *MUST* contain the sequence in fasta format at the end of the file as PROKKA does. `load_gff_list()` parses the files and creates different directories to store all type of files FNA (whole genome sequence), FFN (CDS sequence) and FAA (aminoacid sequence) besides the original GFF files. You can specify what kind of file you want to use in most of the main functions.

The objects created by `load_gff_list()` can be reused in other R sessions.

```r
gff_files <- dir("./examplePATO", pattern = ".gff", full.names = T)
gffs <- load_gff_list(gff_files)
```

**`mash()`** It is a wrapper of Mash. Mash is a method for "Fast genome and metagenome distance estimation using MinHash"(). MASH accept nucleotide or aminoacid fasta files and estimates a similarity distance. To extend the information we recommend to read the main paper *Mash: fast genome and metagenome distance estimation using MinHash. Ondov BD, Treangen TJ, Melsted P, Mallonee AB, Bergman NH, Koren S, Phillippy AM. Genome Biol. 2016 Jun 20;17(1):132. doi: 10.1186/s13059-016-0997-x*

To create a `mash` object you need to supply a list (a vector indeed) of files or a `gff_list` object. By default `mash` use Amino Acids fasta files but you can provide Nucleotide FASTA files (CDS or WGS).

| Input -> I<br>Output -> O | Functions | Objects | | | | | | | | |
|---|---|---|---|---|---|---|---|---|---|---|
| | | files | mmseq | gf _list | core_genome | core_snp_genome | mash | accnet | accnet_enr | igraph |
| QC | classif er | I | | I | | | | | | |
| | extract_non_redundant | | | | | | I/O | I/O | | |
| | non_redundant_hier | | | | | | I | I | | |
| | non_redundant_pangenomes | I | | | | | | | | |
| | non_redundant | | | | I | | I | I | | |
| | outliers | | | | | | I | I | | |
| | remove_outliers | | | | | | I/O | I/O | | |
| Main Functions | load_gf _list | I | | O | | | | | | |
| | accnet | | I | | | | | O | | |
| | mash | I | | I | | | O | | | |
| | mmseqs | I | | I | | | O | | | |
| | core_genome | | I | | O | | | | | |
| | core_snp_genome | I | | I | | O | | | | |
| | pagenomes_from_mmseqs | | I | | | | | O | | |
| | pangenomes_from_f les | | I | | | | | O | | |
| Visualization | core_plots | | I | | | | | | | |
| | similarity_network | | | | | | I | I | | |
| | similarity_tree | | | | | | I | I | | |
| | snps_map | | | | | I | | | | |
| | umap_plot | | | | | | I | I | | |
| | heatmap_of_annotation | | | | | | | | | |
| | network_of_annotation | | | | | | | | | O |
| | plot_knnn_network | | | | | | | | | I |
| Analysis | accnet_enrichment_analysis | | | | | | | I | O | |
| | accnet_with_padj | | | | | | | O | I | |
| | annotate | | I | | | | | | | |
| | clustering | | | | | | I | I | | I |
| | coincidents | | | | | | | I | | |
| | core_snps_matrix | | | | I | I | | | | |
| | export_accnet_aln | | | | | | | I | | |
| | export_core_to_fasta | | | | I | I | | | | |
| | export_to_gephi | | | | | | | I | | I |
| | knnn | | | | | | I | I | | O |
| | sharedness | | I | | | | | I | | |
| | singles | | | | | | | I | | |
| | snps_pairwaise | I | | I | | | | | | |
| | twins | | | | | | | I | | |
| | plamidome | | | I/O | | | | | | |

Figure 1: Table of the PATO functions

```r
my_mash <- mash(gffs, type ="prot")
#or
my_mash <- mash(my_file_list, type ="prot")
```

Besides, you can change some parameters such as `n_cores`, `sketch`, `kmer` and `type`.

```r
my_mash <- mash(gffs, n_cores = 20,
sketch = 1000, kmer = 21, type = "prot")
```

**`mmseqs()`** It is a wrapper of MMSeqs2 software. Creators of MMSeqs define their software as:

> *MMseqs2 (Many-against-Many sequence searching) is a software suite to search and cluster huge protein and nucleotide sequence sets. MMseqs2 is an open source GPL-licensed software implemented in C++ for Linux, MacOS, and (as beta version, via cygwin) Windows. The software is designed to run on multiple cores and servers and exhibits a very good scalability. MMseqs2 can run 10000 times faster than BLAST. At 100 times its speed achieves almost the same sensitivity. It can perform profile searches with the same sensitivity as PSI-BLAST at over 400 times its speed.*

The algorithm renames all the files first, creating a new unique header for each gene/protein. Then the algorithm clusters all the sequences using the `mmseqs linclust` pipeline. Then the clustering results are loaded in R and parse to create the `mmseq` object.

`mmseqs()` can handle a list of files or a `gff_list` object but in this case you only can use feature files, it means genes (FFN) or proteins (FAA).

To create a `mmseq` object you just have to run:

```r
my_mmseq <- mmseqs(my_list)
```

You can change default parameter as `coverage`, `identity` or `evalue` among `n_cores`. Also you can set especifict parameters of clustering (searching for protein families and orthologous). These parameters are `cov_mode` and `cluster_mode`.

```r
my_mmseq <- mmseqs(my_list, coverage = 0.8, identity = 0.8, evalue = 1e-6,
                   n_cores = 20, cov_mode = 0, cluster_mode = 0)
```

The `mmseq` object is key in the analysis of the pangenome definition (core, accessory, pangenome) as well as other process such as annotation or population structure (if you are using accessory to define it). The setting of the parameters can variate the results. Lower values of identity and/or coverage can create larger core-genomes and shorter accessory genomes and vice versa. Besides, the coverage mode and the clustering mode also affect the gene/proteins families definition.

**Coverage Mode** (this section has been extracted from https://github.com/soedinglab/MMseqs2/wiki)

MMseqs2 has three modes to control the sequence length overlap "coverage": (0) bidirectional, (1) target coverage, (2) query coverage and (3) target-in-query length coverage. In the context of cluster or linclust, the query is seen representative sequence and target is a member sequence. The `-cov-mode` flag also automatically sets the `cluster_mode`.

**Clustering modes** All clustering modes transform the alignment results into an undirected graph. In this graph notation, each vertex (i.e. node) represents a sequence, which is connected to other sequences by edges. An edge between a pair of sequences is introduced if the alignment criteria (e.g. `identity`, `coverage` and `evalur`) are fulfilled.

The Greedy Set cover (`cluster-mode = 0`) algorithm is an approximation for the NP-complete optimization problem called set cover.

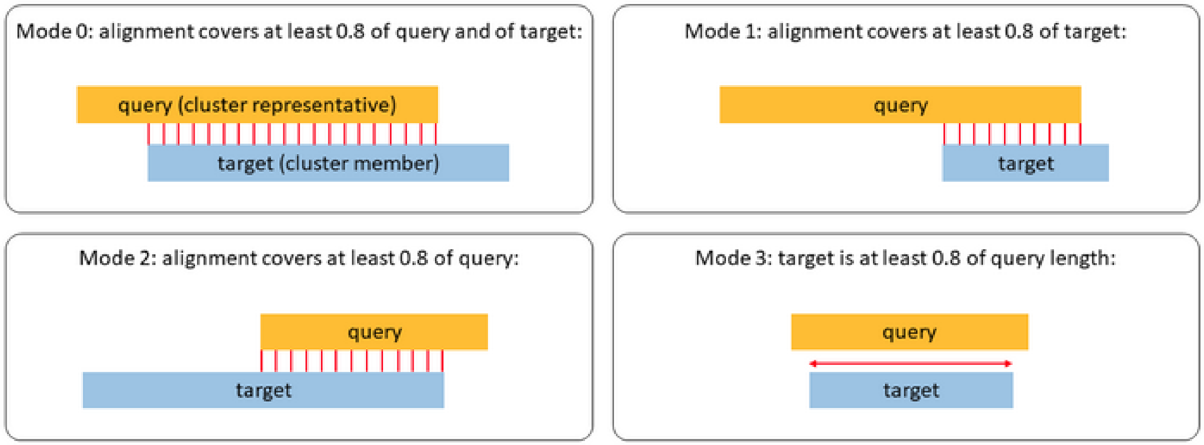

Figure 2: Coverage Mode

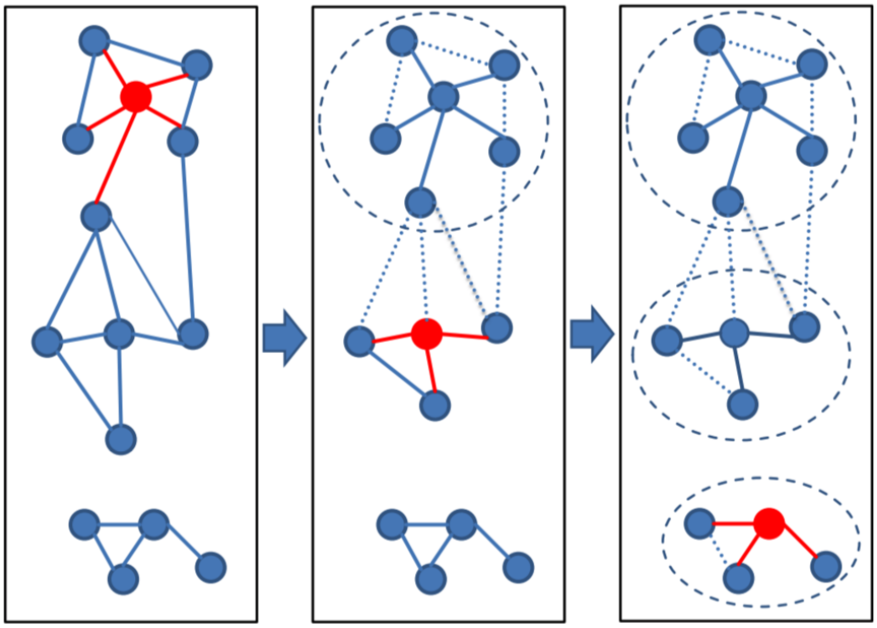

Figure 3: Set Cover clustering

Greedy set cover works by interactively selecting the node with most connections and all its connected nodes to form a cluster and repeating until all nodes are in a cluster.

The greedy set cover is followed by a reassignment step. A cluster member is assigned to another cluster centroid if their alignment score was higher.

Connected component (`cluster_mode = 1`) uses transitive connection to cover more remote homologous.

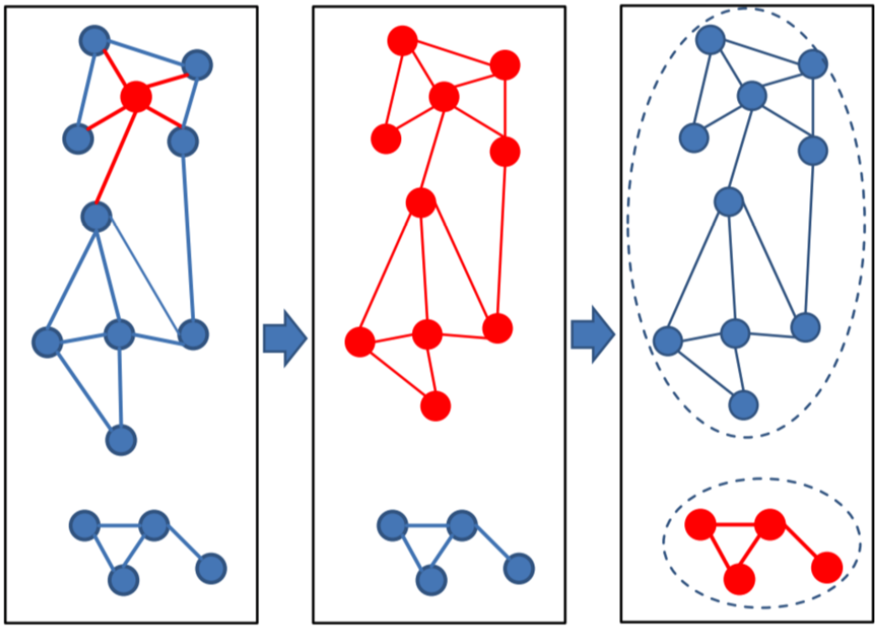

Figure 4: Connected component clustering

In connected component clustering starting at the mostly connected vertex, all vertices that are reachable in a breadth-first search are members of the cluster.

Greedy incremental (`cluster_mode = 2`) works analogous to CD-HIT clustering algorithm.

Greedy incremental clustering takes the longest sequence (indicated by the size of the node) and puts all connected sequences in that cluster, then repeatedly, the longest sequence of the remaining set forms the next cluster.

**`accnet()`** Accnet is a representation of the "accessory genome" using a bipartite network. (see more in *Lanza, V. F., Baquero, F., de la Cruz, F. & Coque, T. M. AcCNET (Accessory Genome Constellation Network): comparative genomics software for accessory genome analysis using bipartite networks. Bioinformatics 33, 283–285 (2017)*). Accnet creates a graph using two kind of nodes: genomes and protein/genes. A genome is connected to a protein/gene if the genome contains this protein (i.e. a protein that belongs to this protein family/cluster). The `accnet` object is one of the main data types in PATO. PATO can create some representations of the accessory genome (dendrograms, networks, plots. . . .) and has some functions to analyze the accessory genome properly. The function `accnet()` builds an `accnet` object taking into account the maximum frequency of each protein/gene to be considered as part of the accessory genome. Moreover, the user can decide to include single protein/genes or not (those which are only present in one sample).

Accnet depends on the definition of protein families, so the input of the `accnet()` function is a `mmseqs` object. The definition of the homologous cluster is critical to define the accessory genome.
`accnet()` can be called as:

```
my_accnet <- accnet(my_mmseq, threshold = 0.8, singles = TRUE)
```

or piping the `mmseqs()` function

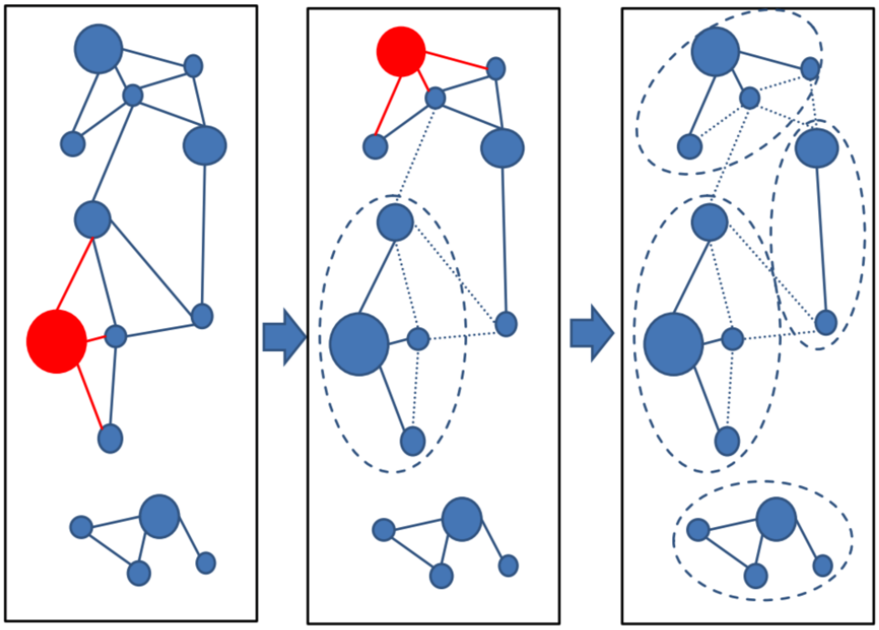

Figure 5: Greedy incremental clustering

```
my_accnet <- mmseqs(my_files) %>% accnet(threshold = 0.8)
```

**pangenomes_from_files()** This function performs clustering using a designated maximum distance among samples and creates a set of pangenomes using it. Pan-genomes can be filtered by genome frequency (i.e. the number of genomes that belong to the pangenome). pangenomes_from_files() builds a new accnet object that relates the pangenome with the genes/proteins. The user can filter the proteins that are included by the presence frequency in the set.

These functions have been designed to work with large datasets (>5.000 genomes). It can be performed as a tailored pipeline to produce an `accnet` object but in a faster way.

**pangenomes_from_mmseqs()** This function creates clusters of genomes (pan-genome) using a list of predefined clusters. The list of clusters can come from an external source (e.g. mlst or BAPS) or from an internal function (e.g clustering function). Moreover, pan-genomes can be filtered by genome frequency (i.e. the number of genomes that belong to the pangenome). `pangenomes_from_mmseqs()` builds a new accnet object that relates the pangenome to the genes/proteins. The user can filter the proteins that are included by the presence frequency. We do not recommend to use it in datasets larger than 5.000 genomes. This function is dependent on an `mmseq` object. The creation of such objects and their size are $O(n^2)$ dependent, so datasets larger than 5000 genomes may exceed computational capabilities.

**core_genome()** It finds and creates a core-genome alignment. Unlike core_plots() this function finds the hard core-genome (genes present in 100% of the genomes and without repetitions). The function takes a `mmseqs()` output, so the definition of the paralogous genes of the core-genome (similarity, coverage and/or e-value) depends on the `mmseqs()` parameters.

The function performs a pseudo-multiple sequence alignment (MSA) per each paralog using the function `result2msa` of mmseqs2 in the proteins case and a mapping with blast (you need to have it installed and in the PATH) if we are using DNA sequences. This approach is much faster than the classical MSA (clustal, mafft or muscle) but it is less accurate. Taking into account that most of the phylogenetic inference software only take variant columns with no insertions or deletions, there are not too many differences in the final phylogenetic trees. `core_genome()` can build a core-genome alignment of thousands of genomes in minutes (see Benchmarking section).

**`core_snp_genome()`** This function finds the core snp genome. Citing Torsten Seemann :

> If you call SNPs for multiple isolates from the same reference, you can produce an alignment of "core SNPs" which can be used to build a high-resolution phylogeny (ignoring possible recombination). A "core site" is a genomic position that is present in all the samples. A core site can have the same nucleotide in every sample ("monomorphic") or some samples can be different ("polymorphic" or "variant"). If we ignore the complications of "ins", "del" variant types, and just use variant sites, these are the "core SNP genome".

This function uses minimap2 to align all the genomes to a reference genome. If reference genome is not specified then `core_snp_genome()` takes one randomly. Once we have all the genomes aligned we look for the conserved regions between all the genomes. Then, we call for the variants using the tool provided with minimap2: paftools.

Finally the SNPs are filtered by the common regions to produce the final core SNP genome.

##### Analysis

**`accnet_enrichment_analysis()`** This function performs a statistical test (Hypergeometric test: phyper) of the genes/proteins. It takes the cluster definition from the cluster table and compute the statistical test assuming the frequency of the gene/protein in the cluster vs the frequency of the gene/protein in the population.

**`accnet_with_padj()`** It creates and improves networks using the adjusted p-value as edge-weight. The network can be exported to gephi (export_to_gephi) for a correct visualization.

**`annotate()`** This function annotates virulence factors and the antibiotic resistance genes of the genomes (files). The annotation is performed using mmseqs2 software and the databases Virulence Factor DataBase and ResFinder. The function can re-use the previous computational steps of `mmseqs` or create a new index database from the files. Re-use option shortens the computational time. This method use the algorithm *search* of mmseqs2 so it only return high identity matchs.

**`clustering()`** This function clusters the genomes using mash data, accnet data or igraph data. The object produced by accnet function, mash function and/or knnn data could be clustered. 'Accnet' objects are clustered using the jaccard distance from presence/absence gene/proteins data. 'Mash' object uses the mash distance values as similarity. 'Igraph' objects could be clustered using the methods availables in igraph.

- for accnet objects:
  - mclust: It performs clustering using Gaussian Finite Mixture Models. It could be combined with d_reduction. This method uses the Mclust package. It has been implemented to find the optimal cluster number
  - upgma: It performs a Hierarchical Clustering using UPGMA algorithm. The n_cluster must be provided
  - ward.D2 It performs a Hierarchical Clustering using Ward algorithm. n_cluster must be provided
  - hdbscan: It performs a Density-based spatial clustering of applications with noise using the DBSCAN package. It finds the optimal number of cluster.
- for mash objects:
  - mclust: It performs clustering using Gaussian Finite Mixture Models. It could be combined with d_reduction. This method uses Mclust pack- age. It has been implemented to find the optimal cluster number

  - upgma: It performs a Hierarchical Clustering using UPGMA algorithm. The n_cluster must be provided
  - ward.D2: It performs a Hierarchical Clustering using the Ward algorithm. The n_cluster must be provided
  - hdbscan: It perform a Density-based spatial clustering of applications with noise using the DBSCAN package. It find the optimal number of cluster.
- for igraph objects
  - greedy: Community structure via greedy optimization of modularity.
  - louvain: This method implements the multi-level modularity optimization algorithm for finding community structure.
  - walktrap: Community structure via short random walks.

Clustering of igraph objects depends on the network building (see knnn function) and the number of cluster may variate between different setting of the k-nn network. Network based-methods are faster than distance based methods.

Dimensional reduction tries to overcome "the curse of dimensionality" (more variables than samples): https://en.wikipedia.org/wiki/Curse_of_dimensionality).) Using `umap` from `uwot` package we reduce to two the dimensionality of the dataset. Note that methods based on HDBSCAN always perform the dimensional reduction. There is not a universal criteria to select the number of clusters and the best configuration for one dataset may not be the best one for others. If you desire to know more about clustering we recommend the book "Practical Guide To Clus-ter Analysis in R" from Alboukadel Kassambara (STHDA ed.)

**`coincidents()`** This function finds groups of coincidents genes/proteins in an `accnet` object. Unlike `accnet_enrichment_analysis()`, `concidents()` does not need to define genome/pangenome clusters. The function performs a multi-dimensional scaling using `umap` and then find clusters with `hdbscan`.

**`core_snps_matrix()`** This function calculates the number of SNPs among samples in core genome and returns a SNP matrix. The functions can provide the results normalized by the alignment length (SNP per Megabase) or the raw number of SNPs. It can also remove the columns with one or more insertion/deletion (only in the case of `core_genome` objects).

**`export_accnet_aln()`** This function writes the presence/absence matrix into a fasta alignment (1 = A, 0 = C) file that can be used with any external software.

**`export_core_to_fasta()`** It exports the `core-genome` and the `core_snp_genome` alignment to a Multi-alignment FASTA file.

**`export_to_gephi()`** This function exports a network (`accnet` or `igraph`) to universal format graphml. Big networks, especially `accnet` networks, cannot be visualized in the R environment. We recommend using external software such as Gephi or Cytoscape. Gephi can manage bigger networks and the ForceAtlas2 layout can arrange `accnet` networks up to 1.000 genomes. If an `accnet` object comes from `accnet_with_padj()` function, the graph files will include the edges-weight proportional to the p-value that associates the protein with the genome.

**`knnn()`** This function creates a network with the k best neighbors of each genome (or pangenome). The network can be built with the best neighbors with or without repetitions. The option `repeats` establish if the best k neighbors includes bi-directional links or not. Let be $G_i$ and $G_j$ two genomes, if $G_i$ is one of the best k-neighbors of $G_j$ and $G_j$ is one of the best k-neighbors of G_i if the repeats option is TRUE then $G_i$ is removed from the k-neighbor and substituted be the next best neighbor if that is not the case G_i is kept in the list.

**plasmidome()** This function uses the **mlplasmids** package to find the contigs belong to plasmids (be aware that **mlplasmdis** only process *Enterococcus faecium*, *Escherichia coli* or *Klebsiella pneumoniae* genomes by now). PATO does not include the **mlplasmids** package by default. It must be installed in your computer manually. The input genomes must be in **gff_list** format (see **load_gff_list()** function). Plasmidome finds the contigs that belong to plasmids and creates a new GFF structure with all the features information (FAA, FNA, FFN and GFF). By default PATO does not install **mlplasmids** because it deposits in its own gitlab repository (https://gitlab.com/sirarredondo/mlplasmids). The authors define mlplasmids as:

> mlplasmids consists on binary classifiers to predict contigs either as plasmid-derived or chromosome-derived. It currently classifies short-read contigs as chromosomes and plasmids for Enterococcus faecium, Escherichia coli and Klebsiella pneumoniae. Further species will be added in the future. The provided classifiers are Support Vector Machines which were previously optimized and selected versus other existing machine-learning techniques (Logistic regression, Random Forest…, see also Reference). The classifiers use pentamer frequencies (n = 1,024) to infer whether a particular contig is plasmid- or chromosome- derived.

To install **mlplasmids** you need to install devtools (which you must have already installed as it is also required for PATO). Then type:

```
devtools::install_git("https://gitlab.com/sirarredondo/mlplasmids")
```

**sharedness()** It creates a matrix with the count of shared genes/proteins among samples.

**singles()** It returns the genes/proteins that are only present in each genome or cluster of genomes.

**snps_pairwaise()** This function calculates the All-vs-All SNPs number. At different of **core_snps_matrix** this function calculates the raw number of SNPs among each pair of sequences avoiding the bias of the core genome size and/or reference. However, this function is an $O(N^2)$ so it can be very computationally expensive for large datasets.

**twins()** Twins nodes are those nodes that share exactly the same edges. In the case of an **accnet** object, twins are those genes/proteins found in the same set of genomes/samples.

##### Quality Control

PATO implements a set of functions to find and remove non desired samples such as other species, outliers or very similar (redundant) samples.

**classifier()** Classifier takes a list of genome files (nucleotide o protein) and identifies the most similar species to each file. **classifier()** uses all reference and representative genomes from NCBI Refseq database and search, using Mash, the best hit for each genome file in the input list.

**extract_non_redundant()** This function creates a non-redundant representation of the input object (**accnet** or **mash**). We recommend using **mash** objects to remove redundancies and redo the **mmseq()** before **accnet()**. Changes in the dataset may affect the clustering process of the genes/proteins and thus the definition of the core-genome.

**non_redundant()** This function creates a non-redundant sub-set of samples. The function accepts an **accnet** or **mash** object and returns a **nr_list** object. The sub-set can be created using a certain distance (mash or jaccard distance depending on the input), a specific number of elements or a fraction of the whole sample set. The function performs an iterative search, so sometimes, the exact number of returned elements could not be the same that the specified in the input. This difference can be defined with the threshold parameter.

**non_redundant_hier()** It creates a non-redundant sub-set of samples. The function accepts an `accnet` or a `mash` object and returns a `nr_list object`. The sub-set can be created using a certain distance (mash or jaccard distance depending on the input), a specific number of elements or a fraction of the whole sample set. The function performs an iterative search so sometimes the exact number of returned elements could not be the same that the specified in the input. This difference can be defined with the threshold parameter. This function performs a hierarchical search. For small datasets (<2000) non_redundant() could be faster. This function consumes less memory than the original so it should be chosen for large datasets.

**non_redundant_pangenomes()** This function removes redundant sequences from a file list of sequences (nucleotide or protein). It has been designed to remove redundant sequences from a dataset. Unlike other non-redundant functions, this function only accepts a distance threshold and has been designed to remove similar sequences in very large datasets (>10.000).

**outliers()** It finds outliers in the dataset using a threshold distance (mash or jaccard distance depending on the input) or by standard deviation times. This function also plots a manhattan plot to visualize the structure of the distance distribution of the input object.

**remove_outliers()** It removes the outliers (or any desired element) of an accnet or mash object.

##### Visualization

One of the reasons to use R as a language to develop PATO is the power of R in terms of visualization. That is why we have implemented several functions to visualize results. Furthermore, most of the PATO objects can be used with all the visualization packages available in R.

**core_plots()** It takes a `mmseq` object and plots the size of core, accessory and pan-genome data in the specified points and with the specified number of replicates. Generally a gene/protein of the core-genome must be present in all the genomes of the dataset. Nevertheless, by random selection and/or errors in sequencing process (sequencing, assembly, ORF finding etc..) some genes/proteins could be missing. For that reason `core_plots()` implements a bootstrapping to create a distribution in each point.

**similarity_network()** The function creates a similarity network connecting genomes with 'distances' lower than the specified threshold. The distance must be suitable for the type of data. MASH values ranging between 0-0.05 intra-specie mean while `accnet` distances could be higher. Unlike `knnn()` each genome is connected with all the genomes closer than the threshold distance.

**similarity_tree()** It builds a similarity tree (pseudo-phylogenetic) of the samples, using whole-genome (`mash`) or accessory genome (`accnet`).

**snps_map()** `snps_map` is a function that makes a similar plot to that of manhattan plots. This function plots the cumulative number of snps per each contig-position pair.

**umap_plot()** UMAP (Uniform Manifold Approximation and Projection for Dimension Reduction) is a dimension reduction technique that can be used for visualisation similarly to t-SNE or PCA, but also for general nonlinear dimension reduction. This function uses the UMAP function implemented in the `uwot` package to represent the population structure of the dataset (`accnet` or `mash`) in 2D plot..

**heatmap_of_annotation()** It creates a heatmap with the `annotate()` function results. The User can filter the results by identity and/or evalue. This function uses `pheatmap` instead of heatmap function if `pheatmap` is installed.

**network_of_annotation()** This function builds an igraph object with the `annotate()` function results. It allows to filter the results by identity and/or Evalue.

**plot_knnn_network()** This function uses the R package **threejs** to draw the k-nnn. You can select the layout algorithm to arrange the network. Moreover, the network can visualize any kind of clustering (internal or external) and color the nodes by cluster. Finally the function returns a list with the network, the layout and the colors (to use with other functions)

### Examples

#### Quality Control and DataSet manipulation

This is an example analyzing 2.941 high quality *Escherichia coli and Shigella* downloaded from NCBI Assembly database (you can download a copy from https://doi.org/10.6084/m9.figshare.13049654). In this case, we will work with protein multi-fasta files (i.e. *.faa).

PATO can handle both nucleotide and aminoacid multi-fasta files or GFF files that include the FASTA sequence (as PROKKA output does). PATO can use whole-genome sequence files but it's restricted for some functions. For small datasets, we recommend to use GFF files which contain all possible formats. For large datasets, we recommend using FAA or FFN files. If you are using NCBI, EBI or other public database note that there are specific files for multi-fasta genes file or multi-fasta protein files. GFF of these database DOES NOT have the FASTA sequence so they cannot be used in `load_gff_list()` function. It is important to know that PATO takes the headers of the fasta sequence as annotation, and usually, protein annotation is better than gene (CDS) annotation. Thus, we recommend to use protein fasta files or reannote the CDS.

It is very important to establish the working directory. We recommend to set the working directory in the folder where the multi-fasta files are. Make sure that you have enough free disk space. Several functions of PATO creates a temporary folder to re-use the data with other functions or sessions. Please do not modify that folder.

```
library(pato)
# We strongly recommend to load tidyverse meta-package
library(tidyverse)
setwd("/myFolder/")
```

The main input file for PATO is a list of files with the path to the multi-fasta files. This list can be easily done by using the `dir()` function. We strongly recommend to use the `full.names = T` parameter to avoid problems with the working directory definition. We have designed PATO to use the base name of the files as input names for the genomes, so you can use the full path in the file lists without fear of affecting the results. You can even join files that are in different folders by simply concatenating the file lists.

```
files <- dir(pattern = ".faa", full.names = T)
```

##### Species and Outliers

PATO has implemented a classification function to identify the most probable species for each input file. The method is based on the minimum `mash` distance of each file against a homemade reference database. The reference database comprises all *reference* and *representative* genomes from Refseq. Input files can be fasta nucleotides or aminoacids.

```
species <- classifier(files,n_cores = 20,type = 'prot')

species %>% group_by(organism_name) %>% summarise(Number = n())

# A tibble: 8 x 2
  organism_name                              Number
  <chr>                                       <int>
1 Citrobacter gillenii                            1
2 Escherichia coli O157:H7 str. Sakai           323
```

```
3 Escherichia coli str. K-12 substr. MG1655   2520
4 Escherichia marmotae                           4
5 Shigella boydii                               18
6 Shigella dysenteriae                           7
7 Shigella flexneri 2a str. 301                 45
8 Shigella sonnei                               40
```

In this example, we found 5 different genomes (1 Citrobacter and 4 E. marmotae). Now we can create a new list of files with the output of the classifier. Using the tidyverse style we can filter out all the unwanted sequences.

```
files = species %>%
  filter(!grepl("Citrobacter",organism_name)) %>%
  filter(!grepl("marmotae",organism_name))
```

Now we create the main objects `mash`and `mmseq`. We establish protein families as 80% of identity, 80% of coverage and a maximum of 1e-6 of E-value.

```
ecoli_mash_all <- mash(files, n_cores = 20, type = 'prot')
ecoli_mm<- mmseqs(files, coverage = 0.8, identity = 0.8, evalue = 1e-6,
                  n_cores = 20)
```

In our case, we use 20 cores to compute the clustering. Most of the PATO functions are multithreading. By default PATO uses all cores minus one for all its multithreading functions.

Finally we can create an `accnet` object:

```
ecoli_accnet_all <- accnet(ecoli_mm,threshold = 0.8, singles = FALSE)
```

One of the things that happens when you download genomes from public databases is that sometimes you can find some *outliers*, genomes that does not belong to the selected species. PATO implements a function to identify outliers and to remove them if it is necessary. In this case, we set a threshold of 0.06 (~94% of ANI) to be considered an outlier.

```
outl <- outliers(ecoli_mash_all,threshold = 0.06)
```

You can remove the outliers from `mash` or `accnet` objects.

```
ecoli_mash_all <-remove_outliers(ecoli_mash_all, outl)
ecoli_accnet_all <-remove_outliers(ecoli_accnet_all, outl)
```

(see ) If you want to remove outliers from `files` object you can do it too.

```
files <-  anti_join(files, outl, by=c("V1"="Source"))
```

This command removes the outliers from the original `files` object. We recommend do it before `mmseqs` command.

##### Redundancies

Redundancies due to strain oversampling (e.g. strains causing outbreaks or abundant/overrepresented bacterial lineages) are extremely frequent in databases and preclude accurate analysis of microbial diversity. PATO implements a function to select a non-redundant subset based on a fix distance (similarity), a % of samples or a selected number of genomes. It is an iterative approach so we implement a threshold of tolerance due to some time finding the exact solution is impossible. This function accepts a `mash` or `accnet` object as input. Besides, there is a function to create a new `mash` or `accnet` object with the non-redundant set of genomes. of genomes

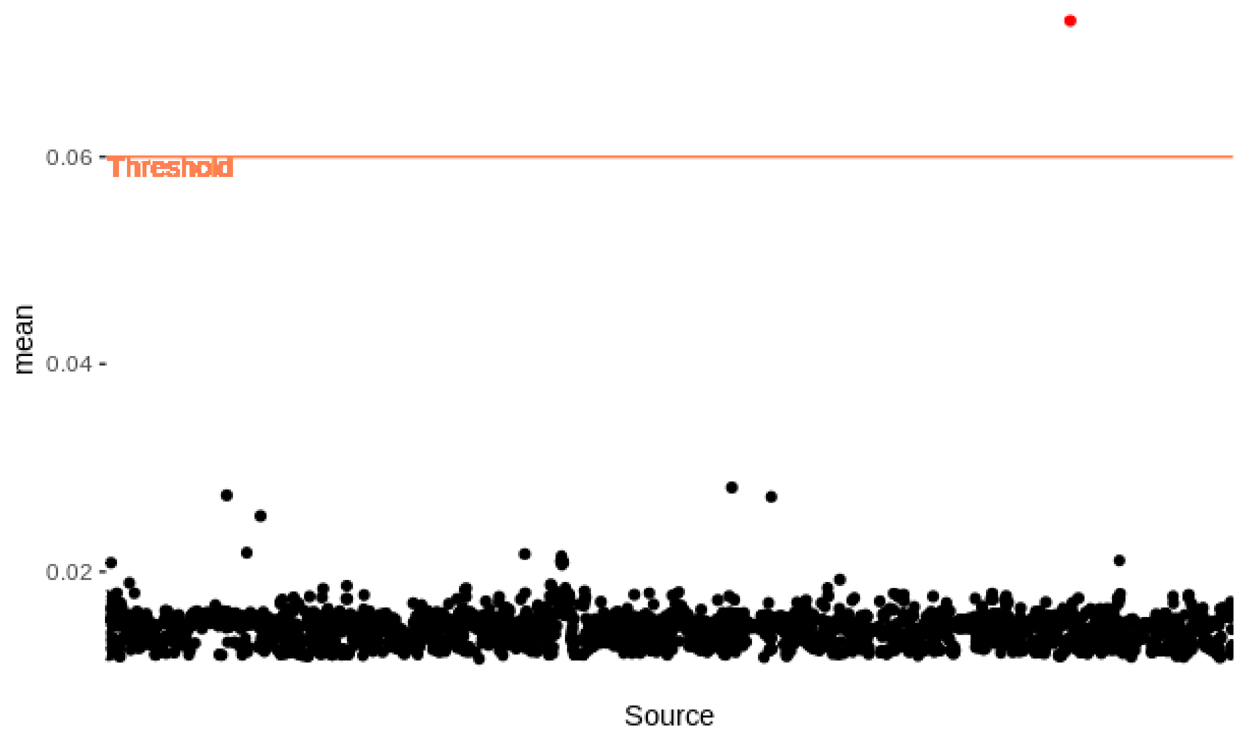

Figure 6: Outliers plot

```
# To select 800 non redundant samples
nr_list <- non_redundant(ecoli_mash_all,number = 800)
```

or

```
# To select a subset of samples with 99.99% of identity
nr_list <- non_redundant(ecoli_mash_all, distance = 0.0001)
```

to create the objects only with the representatives of each cluster:

```
ecoli_accnet <- extract_non_redundant(ecoli_accnet_all, nr_list)
ecoli_mash <- extract_non_redundant(ecoli_mash_all, nr_list)
```

Moreover, there is another function `non_redundant_hier()` that performs a hierarchical approach to cluster the genomes. It is faster for a big dataset (>1000 genomes) so we recommend using it in these cases.

```
nr_list <- non_redundant_hier(ecoli_mash_all,800, partitions = 10)
ecoli_accnet <- extract_non_redundant(ecoli_accnet_all, nr_list)
ecoli_mash <- extract_non_redundant(ecoli_mash_all, nr_list)
```

The function uses a "divide and conquer" approach. It divides the total set into $n$ partitions and groups them separately. Finally, it re-clusters the results for each of the partitions. This operation is repeated in each iteration until the required number of terms is reached.

#### Population Structure / Pangenome Description

In this example we continue with the *Escherichia coli and Shigella* high quality dataset. You can download a copy from https://doi.org/10.6084/m9.figshare.13049654). We continue after the QC described above.

##### Core Genome analysis

PATO implements a set of tools to inspect the core genome. It includes the pangenome composition (core, accessory and pangenome) and the function to create a core-genome alignment or core-snp-genome..

We can inspect the Pangenome composition of our dataset with:

```
cp <- core_plots(ecoli_mm,reps = 10, threshold = 0.95, steps = 10)
```

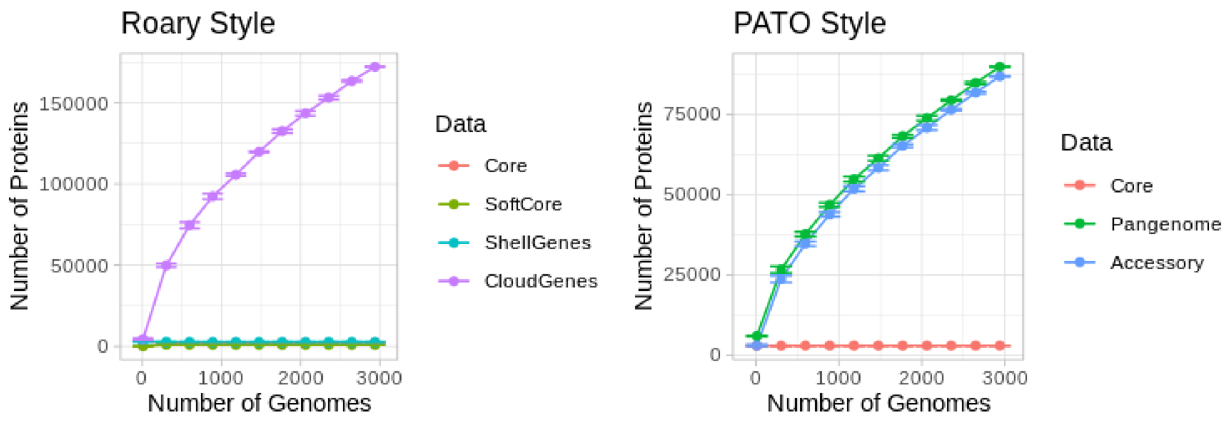

Figure 7: Plots of the pangenome composition

PATO can build a core-genome alignment to use with external software such as IQTree[http://www.iqtree.org/], RAxML-NG[http://www.exelixis-lab.org/software.html], FastTree[http://www.microbesonline.org/fasttree/] or other phylogenetic inference software. The core genome is computed based on a `mmseq` object, so the definition of the core-genome depends on the parameters used in that step.

In this example we build a core-genome alignment of the ~800 samples from non-redundant result.

```
nr_files = nr_list %>%
  as.data.frame() %>%
  group_by(cluster) %>%
  top_n(1,centrality) %>%
  summarise_all(first) %>%
  select(Source) %>%
  distinct()
```

We have selected the best sample of each cluster (max centrality value). Some cluster has several samples with the same centrality value, so we take one of them (the first one). Now we run the core genome for this list of samples.

```
core <- mmseqs(nr_files) %>% core_genome(type = "prot")
export_core_to_fasta(core,"core.aln")
```

Sometimes, when you are using public data, core-genome can be smaller than you would expect. Citing **Andrew Page**, creator of Roary (https://sanger-pathogens.github.io/Roary/)

> ***I downloaded a load of random assemblies from GenBank. Why am I seeing crazy results?***
> *Gene prediction software rarely completely agrees, with differing opinions on the actual start of a gene, or which of the overlapping open reading frames is actually the correct one, etc. As a result, if you mix these together you can get crazy results. The solution is to reannotate all of the genomes with a single method (like PROKKA). Otherwise you will waste vast amounts of time looking at noise, errors, and the batch effect.*

Other times you can have a problem of *outliers*

The exported files are in Multi-alignment FASTA format and ca be use with most of the Phylogenetic tools. In this case we used FastTree for phylogenetic inference

```
fasttreeMP core.aln > core.tree
```

And then, we can read the output file to import to our R environment and plot the result.

```
library(phangorn)
library(ggtree)
core_tree = read.tree("core.tree")

annot_tree = species %>%
  filter(!grepl("Citrobacter",organism_name)) %>%
  filter(!grepl("marmotae",organism_name)) %>%
  select(Target,organism_name) %>% distinct()
core_tree %>% midpoint %>% ggtree(layout = "circular") %<+%
  annot_tree + geom_tippoint(aes(color = organism_name))
```

In this case, we have added the species information extracted above.

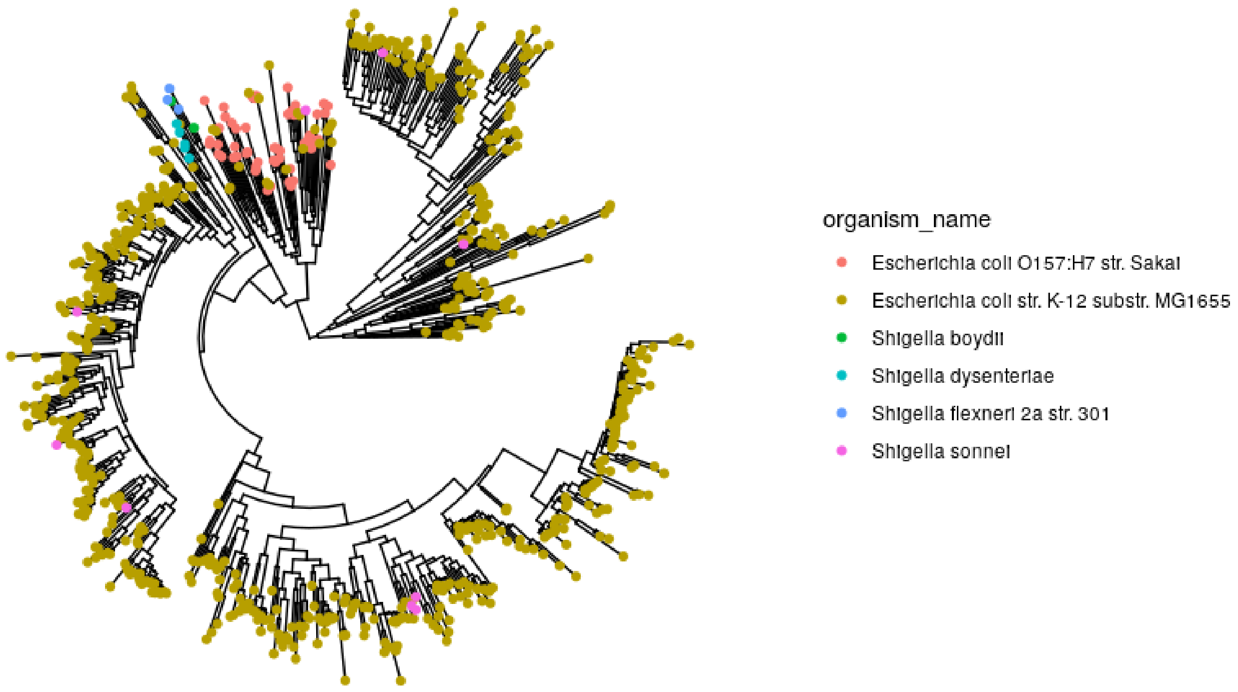

Figure 8: ML Core-genome phylogenetic tree

You must take into account that the Maximum Likelihood tree can take long computational times.

Finally, you can obtain a SNPs matrix from the core genome alignment. The matrix shows the number of total SNPs (or changes in the case of proteins alignments) shared among the samples. The matrix can be normalized by the total length of the genome (in magabases) or can be a raw number of variants.

```
var_matrix = core_snps_matrix(core, norm = TRUE)
pheatmap::pheatmap(var_matrix,show_rownames = F, show_colnames = F)
```

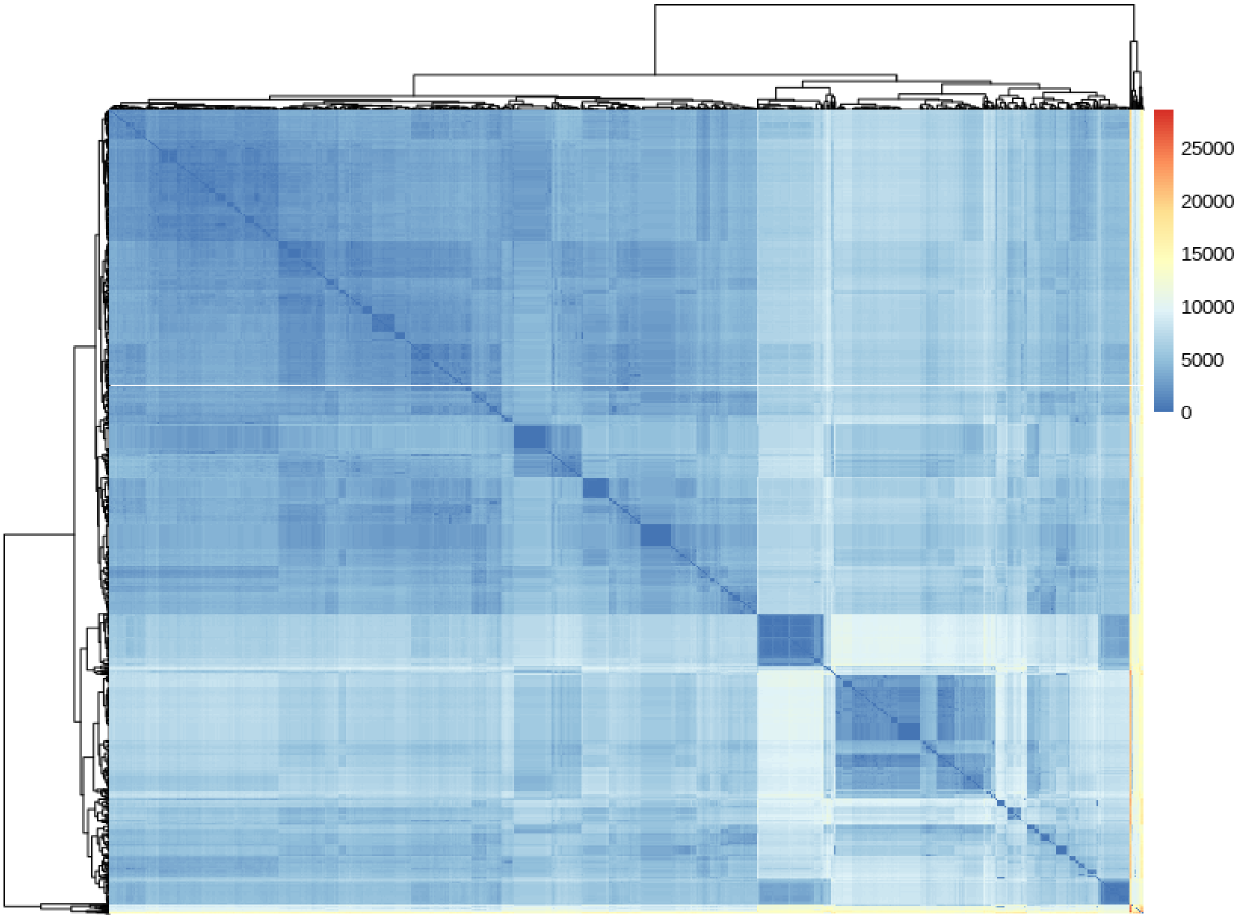

Figure 9: HeatMap of the *core-snp-genome* SNP matrix

##### Accessory Genome Analysis

I this case we are creating the *accessory genome* taking those proteins present in no more than 80% of the genomes.

```
ecoli_accnet_all <- accnet(ecoli_mm,threshold = 0.8, singles = FALSE)
```

As we have shown above, PATO has an object type of the accessory genome `accnet` and functions to analyze and visualize the content of the accessory genome and the relationship between genes/proteins and the genomes. We can visualize an `accnet` object as a bipartite network. Commonly, AccNET networks are very large, so, we recommend visualizing the networks using Gephi. We do not recommend to try visualizing AccNET networks with more than 1.000 genomes.

One of the most interesting things when we analyze an accessory genome, is try to find what genes/proteins are over-represented in a set of genomes. `accnet_enrichment_analysis()` function analyze what genes/proteins are over-represented in a *cluster* in comparison with the population (i.e. in the whole dataset). The clusters definition can be external (any categorical metadata such as Sequence Type, Source, Serotype etc…) or internal using some clustering process. We have to take into account, that the redundancies can bias this kind of analysis. If there is an over-representation of strains/genomes in your dataset (e.g. genomes from identícal isolates causing an outbreak), the results could result in a biased analysis. You will find more significant genes/proteins because the diversity of the dataset is not homogeneously distributed. For this reason we recommend to use a non redundant set of samples. In this example we select 800 non redundant genomes.

```
ec_nr<- non_redundant(ecoli_mash,number = 800 )
ecoli_accnet_nr <- extract_non_redundant(ecoli_accnet, ef_nr)
ec_800_cl <- clustering(ecoli_accnet_nr, method = "mclust", d_reduction = TRUE)
```

Now we can visualize the network using Gephi.

```
export_to_gephi(ecoli_accnet_nr, "accnet800", cluster = ec_800_cl)
```

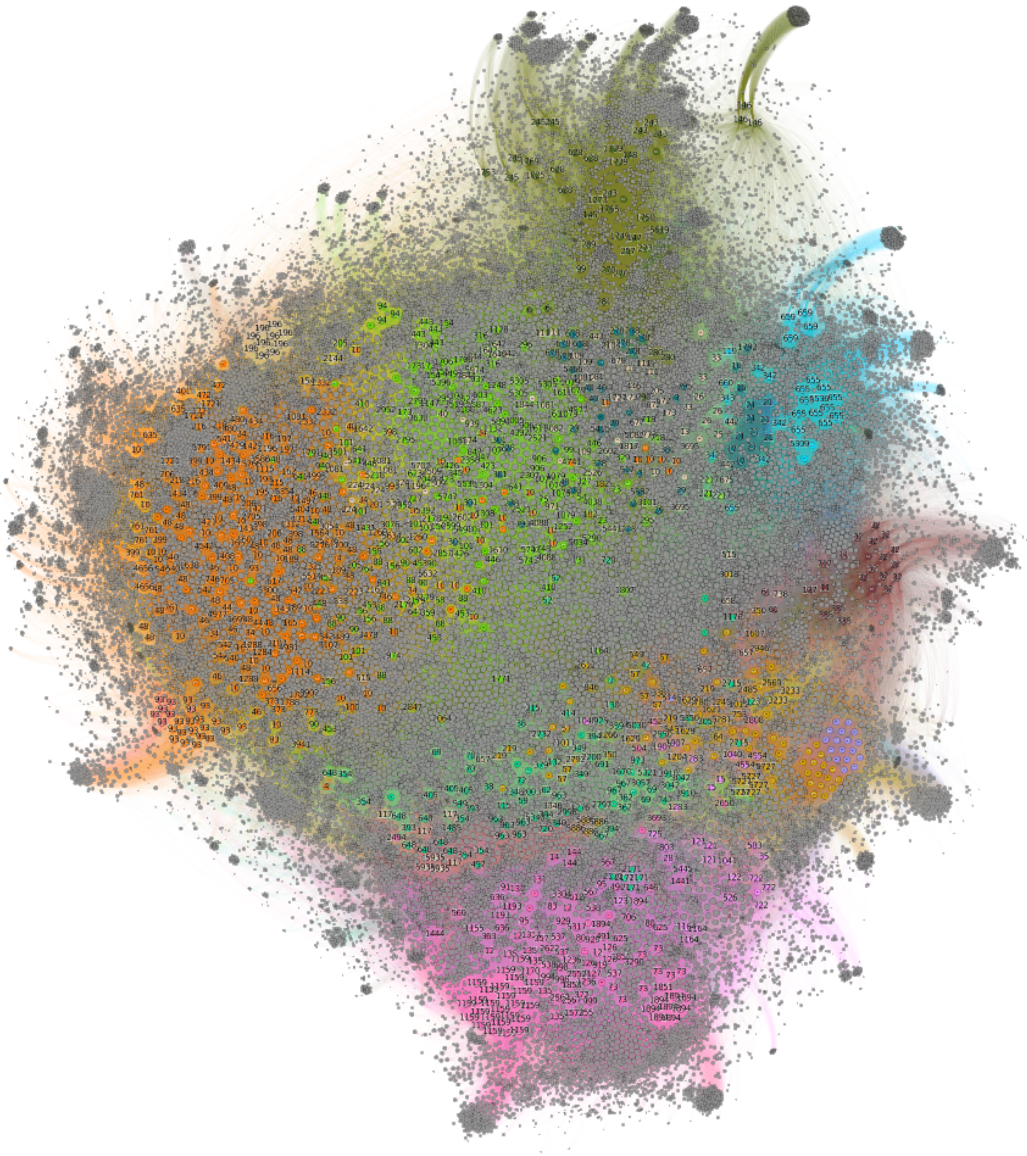

[https://doi.org/10.6084/m9.figshare.13089338]

To perform the enrichment analysis we use:

```
accnet_enr_result <- accnet_enrichment_analysis(ecoli_accnet_nr,
                                                cluster = ef_800_cl)

# A tibble: 1,149,041 x 14
   Target            Source                  Cluster perClusterFreq ClusterFreq
   ClusterGenomeSize perTotalFreq TotalFreq OdsRatio   pvalue    padj
   AccnetGenomeSize AccnetProteinSi... Annot
   <chr>             <chr>                     <dbl>          <dbl>
     <int>             <int>         <dbl>     <int>    <dbl>    <dbl>   <dbl>
     <int>            <int> <chr>
 1 1016|NZ_CP053592.... GCF_000009565.1_ASM956v...      1          0.0436
```

```
  13               298       0.0173        20     2.52  3.62e-5 0.00130
  1156            74671 ""
 2 1016|NZ_CP053592.... GCF_000022665.1_ASM2266...       1         0.0436
 13               298       0.0173        20     2.52  3.62e-5 0.00130
 1156            74671 ""
 3 1016|NZ_CP053592.... GCF_000023665.1_ASM2366...       1         0.0436
 13               298       0.0173        20     2.52  3.62e-5 0.00130
 1156            74671 ""
 4 1016|NZ_CP053592.... GCF_000830035.1_ASM8300...       1         0.0436
 13               298       0.0173        20     2.52  3.62e-5 0.00130
 1156            74671 ""
 5 1016|NZ_CP053592.... GCF_000833145.1_ASM8331...       1         0.0436
 13               298       0.0173        20     2.52  3.62e-5 0.00130
 1156            74671 ""
 6 1016|NZ_CP053592.... GCF_001039415.1_ASM1039...       1         0.0436
 13               298       0.0173        20     2.52  3.62e-5 0.00130
 1156            74671 ""
 7 1016|NZ_CP053592.... GCF_001596115.1_ASM1596...       1         0.0436
 13               298       0.0173        20     2.52  3.62e-5 0.00130
 1156            74671 ""
 8 1016|NZ_CP053592.... GCF_009663855.1_ASM9663...       1         0.0436
 13               298       0.0173        20     2.52  3.62e-5 0.00130
 1156            74671 ""
 9 1016|NZ_CP053592.... GCF_009832985.1_ASM9832...       1         0.0436
 13               298       0.0173        20     2.52  3.62e-5 0.00130
 1156            74671 ""
10 1016|NZ_CP053592.... GCF_013166955.1_ASM1316...       1         0.0436
13               298       0.0173        20     2.52  3.62e-5 0.00130
1156            74671 ""
# ... with 1,149,031 more rows
```

[https://doi.org/10.6084/m9.figshare.13089314.v2]

Now, we can export a new network with the *adjusted p-values* as edge-weight.

```
accnet_with_padj(accnet_enr_result) %>%
  export_to_gephi("accnet800.padj", cluster = ec_800_cl)
```

[https://doi.org/10.6084/m9.figshare.13089401.v1]

PATO also includes some functions to study the genes/proteins distribution: `singles()`, `twins()` and `conicidents()`.

- `singles()` finds those genes/proteins that are only present in a sample (genome, pangenome…).
- `twins()` finds those genes/proteins that have the same connections (i.e. genes/proteins present in the same genomes).
- `coincidents()` finds those genes/proteins with similar connections (i.e. genes/proteins that usually are together)

The differences between the`twins()` and the `concidents()` are i) the higher flexibility and thus, lower sensitivity to outliers of the `coincidents()`, ii) the higher speed, accuracy and sensitivity to noise or outliers of the twins due to one missed connection (fe.g. a bad prediction for a protein in a genome) removes automatically such protein from the `twin` group. Furtheremore, the low speed of the`coincidents()` sometimes results in big clusters of pseudo-cores.

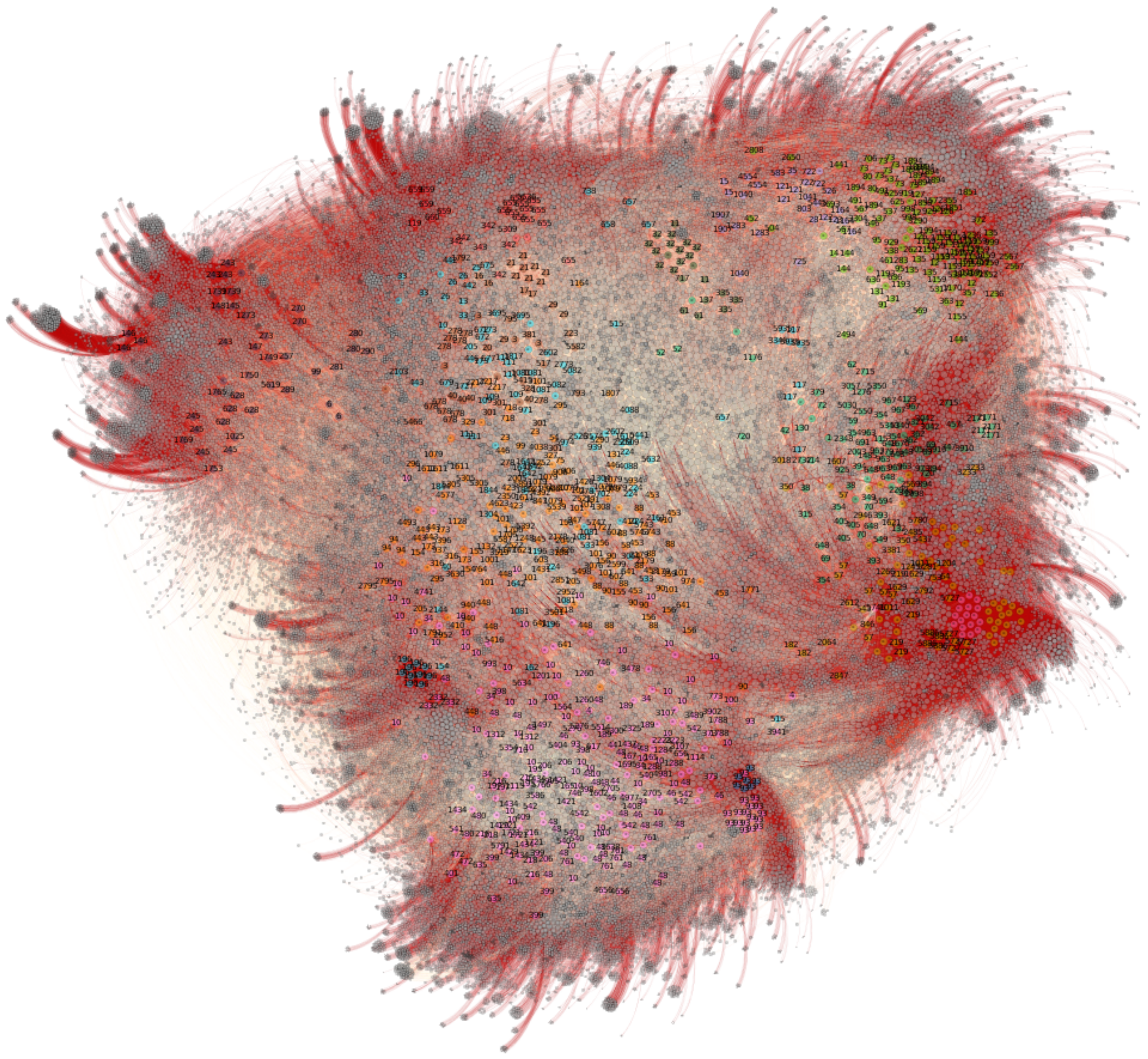

Figure 10: Accnet network with enrichment analysis add. Edge color are proportional to the p-value (adjusted)

##### Population structure

PATO includes tools for population structure analysis n. PATO can analyze the population structure of the whole-genome (MASH based) or the accessory structure (AccNET based)

We can visualize our dataset as a dendrogram (i.e a tree). PATO allows to visualize both `mash` and `accnet` data as a tree. In the case of `accnet` data, first PATO calculates a distance matrix using the presence/absence matrix in combination with Jaccard distance. The function `similarity_tree()` implements different methods to build the dendrogram. Phylogenetic aproaches such as Neighbor Joining, FastME: Minimum Evolution and hierarchical clustering method such as complete linkage, UPGMA, Ward's mininum variance, and WPGMC.

```
mash_tree_fastme <- similarity_tree(ecoli_mash)
mash_tree_NJ <- similarity_tree(ecoli_mash, method = "NJ")
mash_tree_upgma <- similarity_tree(ecoli_mash,method = "UPGMA")
accnet_tree_upgma <- similarity_tree(ecoli_accnet,method = "UPGMA")
```

The output has a `phylo` format, so can be visualized with external packages as `ggtree`.

```
mash_tree_fastme %>% midpoint %>% ggtree(layout = "circular") %<+%
  annot_tree + geom_tippoint(aes(color = organism_name))
```

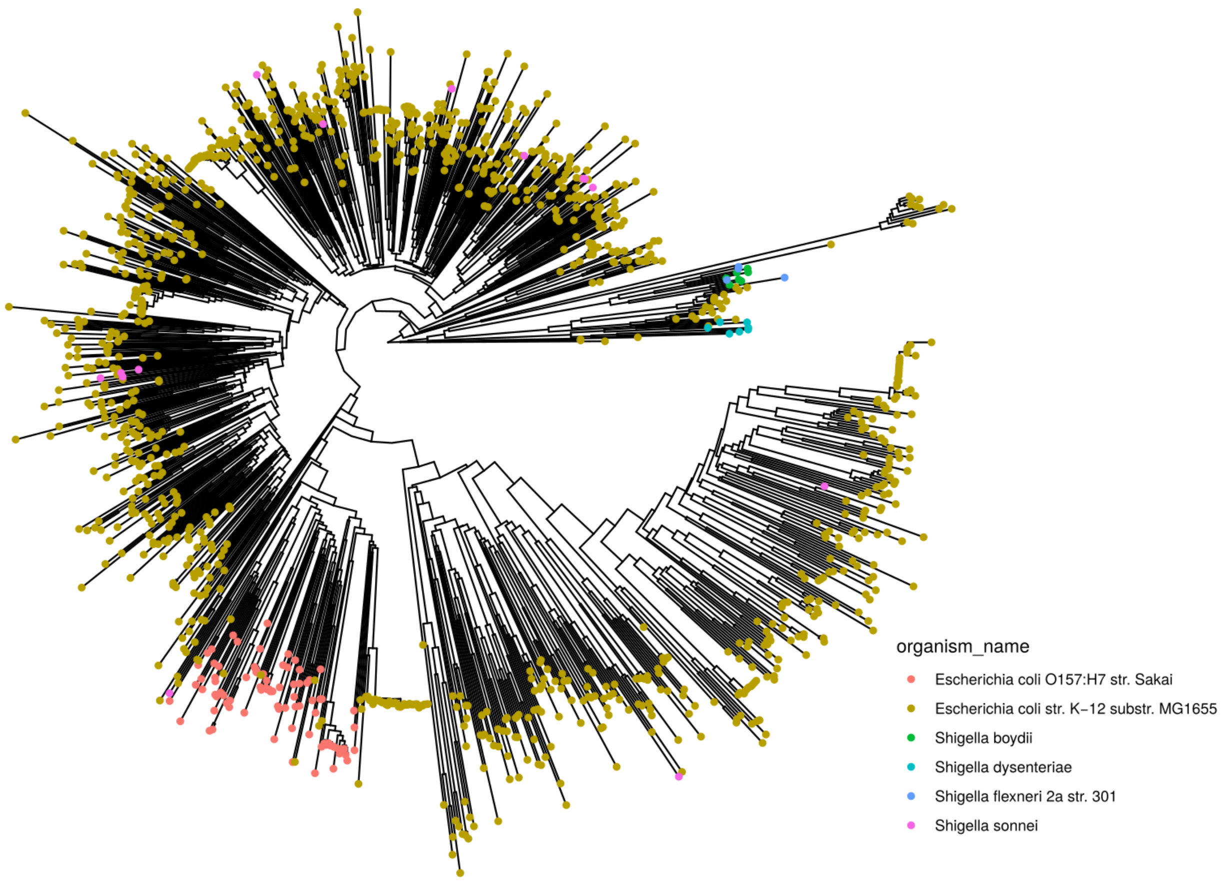

Figure 11: Neighbor Joining tree of MASH distances

Using other external packages, we can compare the arrangement of the pangenome (`mash` data) against the accessory genome (`accnet`)

```
library(dendextend)
tanglegram(ladderize(mash_tree_upgma), ladderize(accnet_tree_upgma),
           fast = TRUE, main = "Mash vs AccNET UPGMA")
```

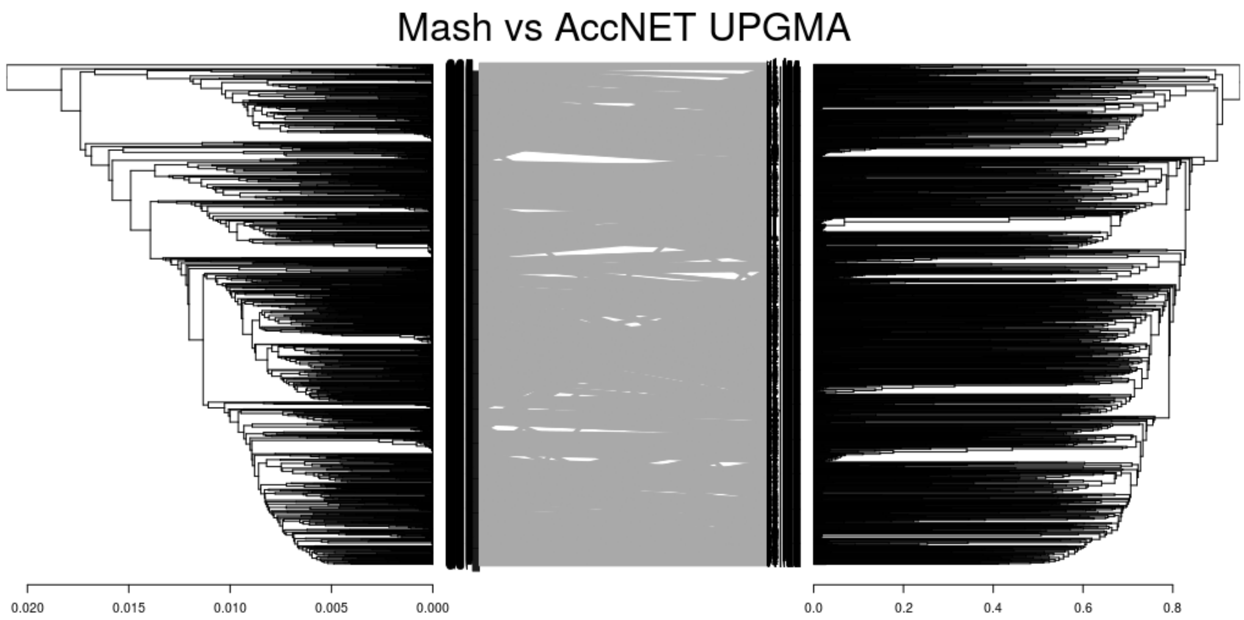

Figure 12: Tanglegram between Mash and AccNET dendrograms

Some Maximum Likelihood inference trees software accept, as input, binary data (0-1) alignments. So, we can use accessory data (*accnet*) to infer a tree (non-phylogenetic) with this data. This is similar to the `similarity_tree()` but instead of being based on distance metrics it's based on ML principles. To export this alignment you can use `export_accnet_aln()`.

```
mmseqs(files) %>% accnet() %>% export_accnet_aln(.,file ="my_accnet_aln.fasta")
```

And then you can use it as input alignment

```
iqtree -s my_accnet_aln.fasta -st BIN
```

Again, we can import to R and plot it.

```
acc_tree = read.tree("acc.aln.treefile")
acc_tree %>% midpoint.root() %>% ggtree()
```

This kind of alignment can be very large. Most of the times, accessory genomes contains a lot of spurious genes/proteins that do not add any information to the alignment. For this reason, `export_accnet_aln()` has a parameter, `min_freq`, to filter the genes/proteins by their frequency. This option cuts significantly the alignment length and improves computational times.

PATO has a set of functions to visualize data. We have seen the trees functions, but it also implements methods to visualize the relationships among genomes as networks. **K-Nearest Neighbor Networks** is a representation of the relationship of the data in a network plot.

```
# K-NNN with 10 neighbors and with repetitions
knnn_mash_10_w_r <- knnn(ecoli_mash,n=10, repeats = TRUE)
# K-NNN with 25 neighbors and with repetitions
knnn_mash_25_w_r <- knnn(ecoli_mash,n=25, repeats = TRUE)
```

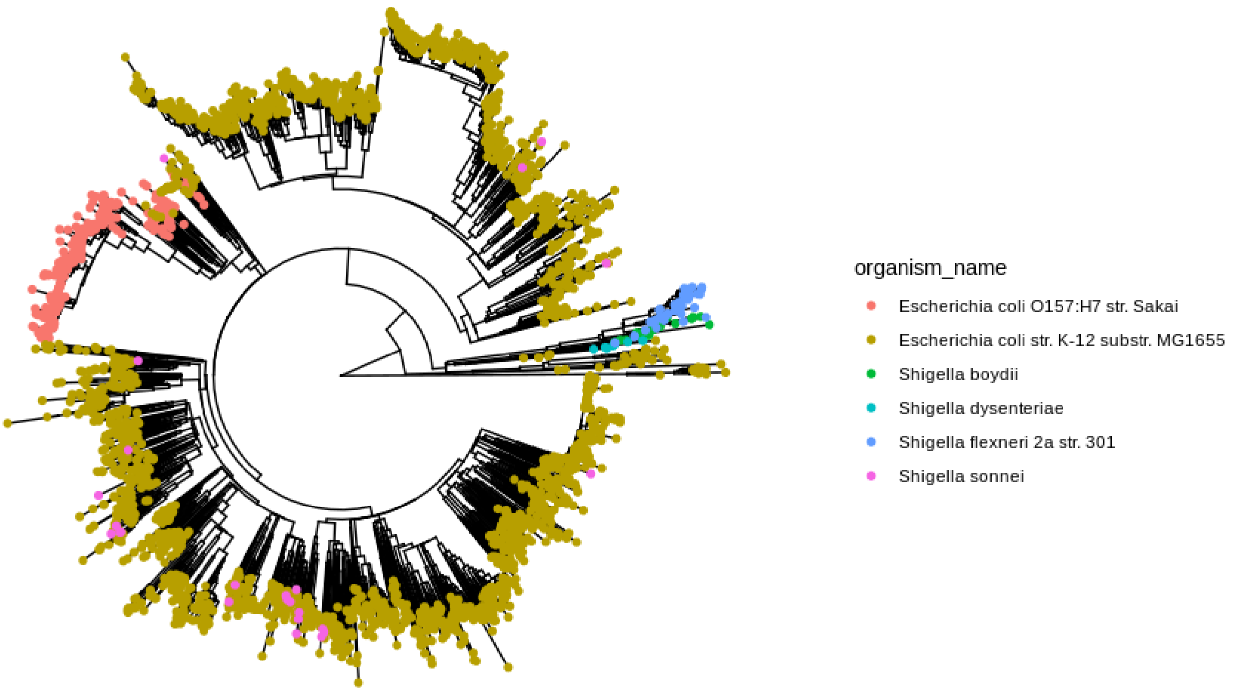

Figure 13: ML tree of accessory genome

```
# K-NNN with 50 neighbors and with repetitions
knnn_mash_50_w_r <- knnn(ecoli_mash,n=50, repeats = TRUE)
```

The `knnn()` function returns an `igraph` object that can be visualized with several packages. However, PATO implements its own function `plot_knnn_network()` that uses `threejs` and its own `igraph` for layouting and plotting the network. Due to the large size of the network we strongly recommend to use this function or use the external software Gephi (https://gephi.org/). You can use the function `export_to_gephi()` to export these networks to Gephi.

```
export_to_gephi(knnn_mash_50_w_r,file = "knnn_50_w_r.tsv")
```

You can use the internal function `plot_knnn_network()` to visualize the network. This function uses `igraph` layouts algorithm to arrange the network and `threejs` package to draw and explore the network.

```
plot_knnn_network(knnn_mash_50_w_r)
```

PATO includes different clustering methods for different kind of data (objects). `clustering()` function joins all the different approaches to make easier the process. For the `mash` and the `accnet`objects we have the following methods:

- **mclust**: It performs clustering using Gaussian Finite Mixture Models. It could be combined with d_reduction. This method uses the Mclust package. It has been implemented to find the optimal cluster number
- **upgma**: It performs a Hierarchical Clustering using the UPGMA algorithm. The user must provide the number of clusters.
- **ward.D2**: It performs a Hierarchical Clustering using Ward algorithm. The user must provide the number of clusters.
- **hdbscan**: It performs a Density-based spatial clustering of applications with noise using the DBSCAN package. It finds the optimal number of clusters.

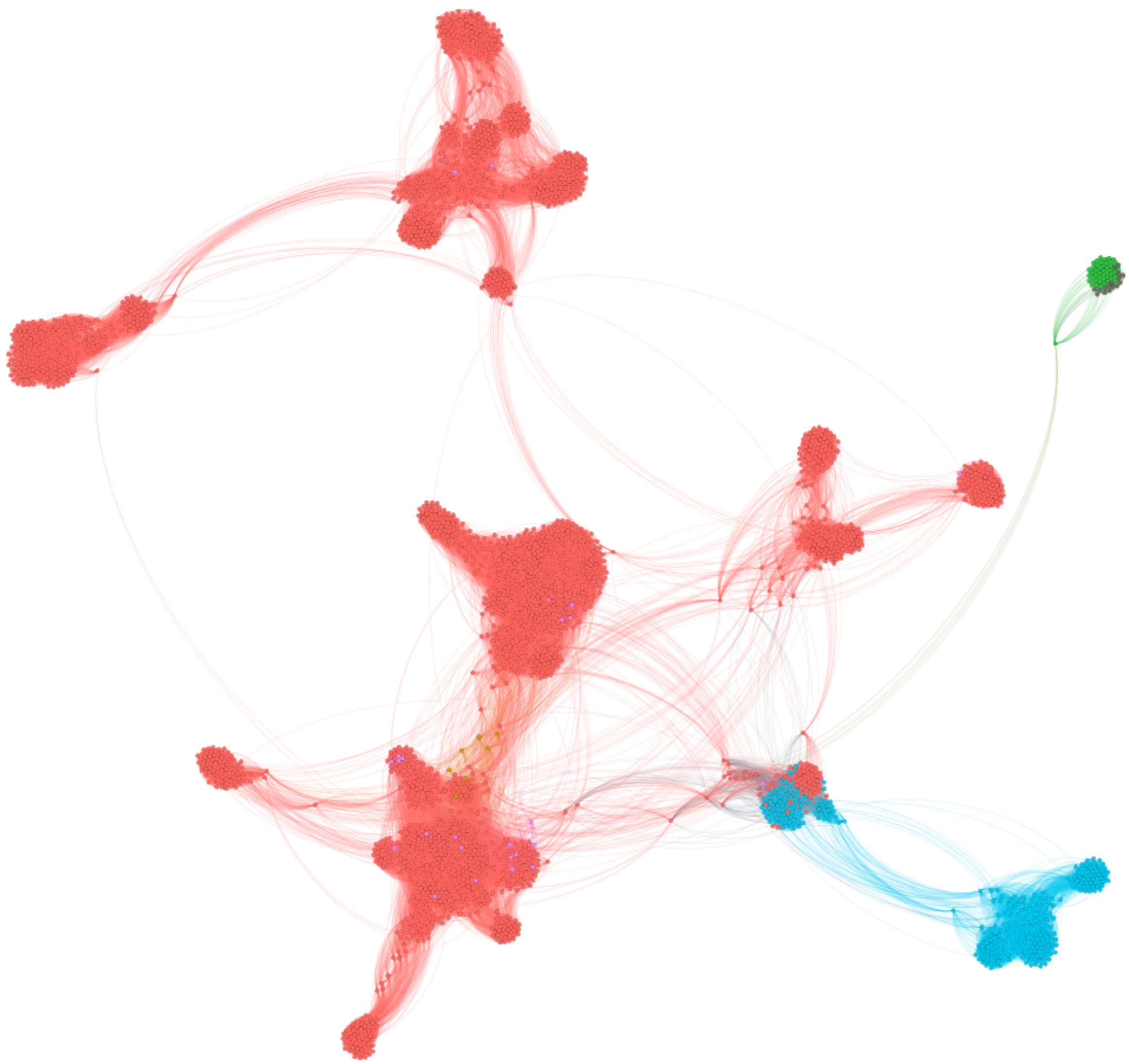

Figure 14: Gephi visualization of the K-NNN with 50 neighbors

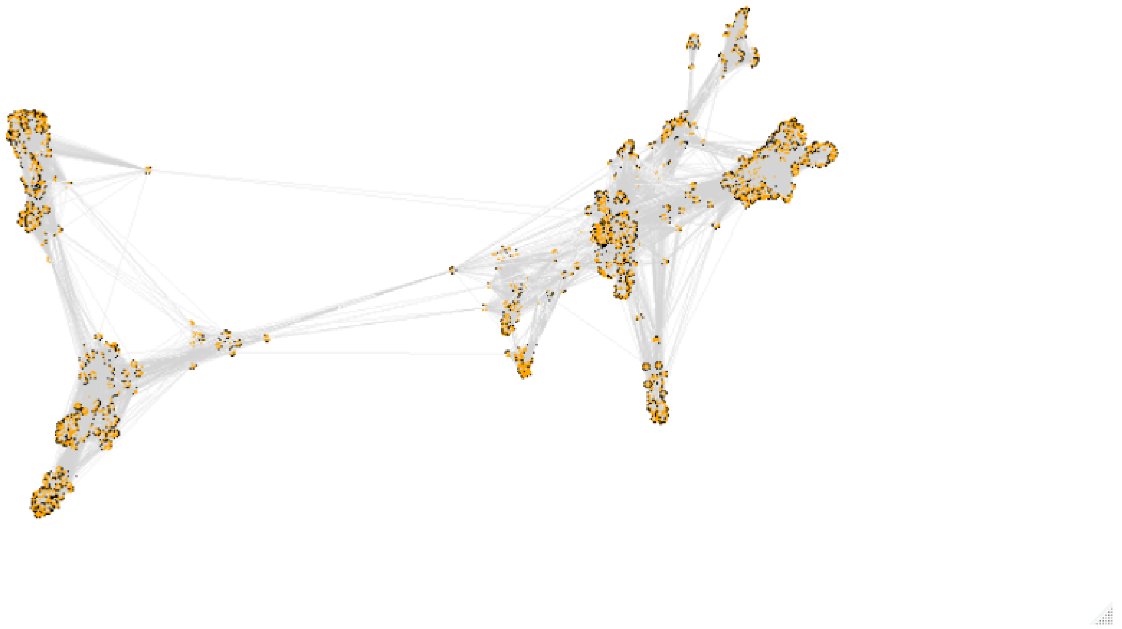

Figure 15: Threejs visualization of the K-NNN with 50 neighbors

Any of the above methods is compatible with a multidimensional scaling (MDS). PATO performs the MDS using **UMAP** algorithm (Uniform Manifold Approximation and Projection). UMAP is a tools for MDS, similar to other machine learning MDS techniques such as t-SNE or Isomap. Dimension reduction algorithms tend to fall into two categories those that seek to preserve the distance structure within the data and those that favors the preservation of local distances over global distance. Algorithms such as PCA , MDS , and Sammon mapping fall into the former category while t-SNE, Isomap or UMAP, fall into the later category. This kind of MDS combine better with the clustering algorithms since clustering process always tries to find local structures.

```
ec_cl_mclust_umap <- clustering(ecoli_mash, method = "mclust",
                                d_reduction = TRUE)
```

Moreover, `clustering()` can handle `knnn` networks as input. In this case , PATO uses network clustering algorithms such as:

- greedy: Community structure via greedy optimization of modularity
- louvain: This method implements the multi-level modularity optimization algorithm for finding community structure
- walktrap: Community structure via short random walks

```
ec_cl_knnn <-clustering(knnn_mash_50_w_r, method = "louvain")
```

Whichever the method used, the output always has the same structure: a `data.frame` with two columns `c("Source","Cluster")`. The reason for this format is to be compatible with the rest of the data, being able to combine with the rest of the objects using the `Source` variable as key.

To visualize the clustering data or just to see the data structure we can use `umap_plot()` function. The functions performs a `umap` reduction and plot the results. We can include the `clustering` results as a parameter and visualize it.

```
umap_mash <- umap_plot(ecoli_mash)
```

```
umap_mash <- umap_plot(ecoli_mash, cluster = ef_cl_mclust_umap)
```

We can also use the `plot_knnn_network()` to visualize the network clustering results.

```
cl_louvain = clustering(knnn_mash_25_wo_r, method = "louvain")
plot_knnn_network(knnn_mash_25_wo_r, cluster = cl_louvain, edge.alpha = 0.1)
```

##### Annotation

PATO has a function to annotate the Antibiotic Resistance Genes and the Virulence Factor: `annotate()`. Antibiotic resistance is predicted using MMSeqs2 over ResFinder database (https://cge.cbs.dtu.dk/services/ResFinder/) (doi: 10.1093/jac/dks261) and Virulence Factors using VFDB (http://www.mgc.ac.cn/cgi-bin/VFs/v5/main.cgi)(doi: 10.1093/nar/gky1080). VFBD has two sets of genes `VF_A` the *core-dataset* and `VF_B` the *full dataset*. The *Core dataset* only includes genes associated with experimentally verified VFs, whereas the *full dataset* includes the genes of the *Core dataset* plus genes predicted as VFs.

The `annotate()` results is a table with all positive hits of each gene/protein of the dataset (files). The `annotate()` table? can re-use the results of `mmseqs()` to accelerate the process. In the same way, the query can be *all* genes/proteins or *accessory* genes/proteins. The results are quite raw so the user must curate the table. We recommend using the `tidyverse` tools.

```
library(tidyverse)

annotation <- annotate(files, type = "prot",database = c("AbR","VF_A"))

annotation %>%
  #remove all hits with identity lower than 95%
```

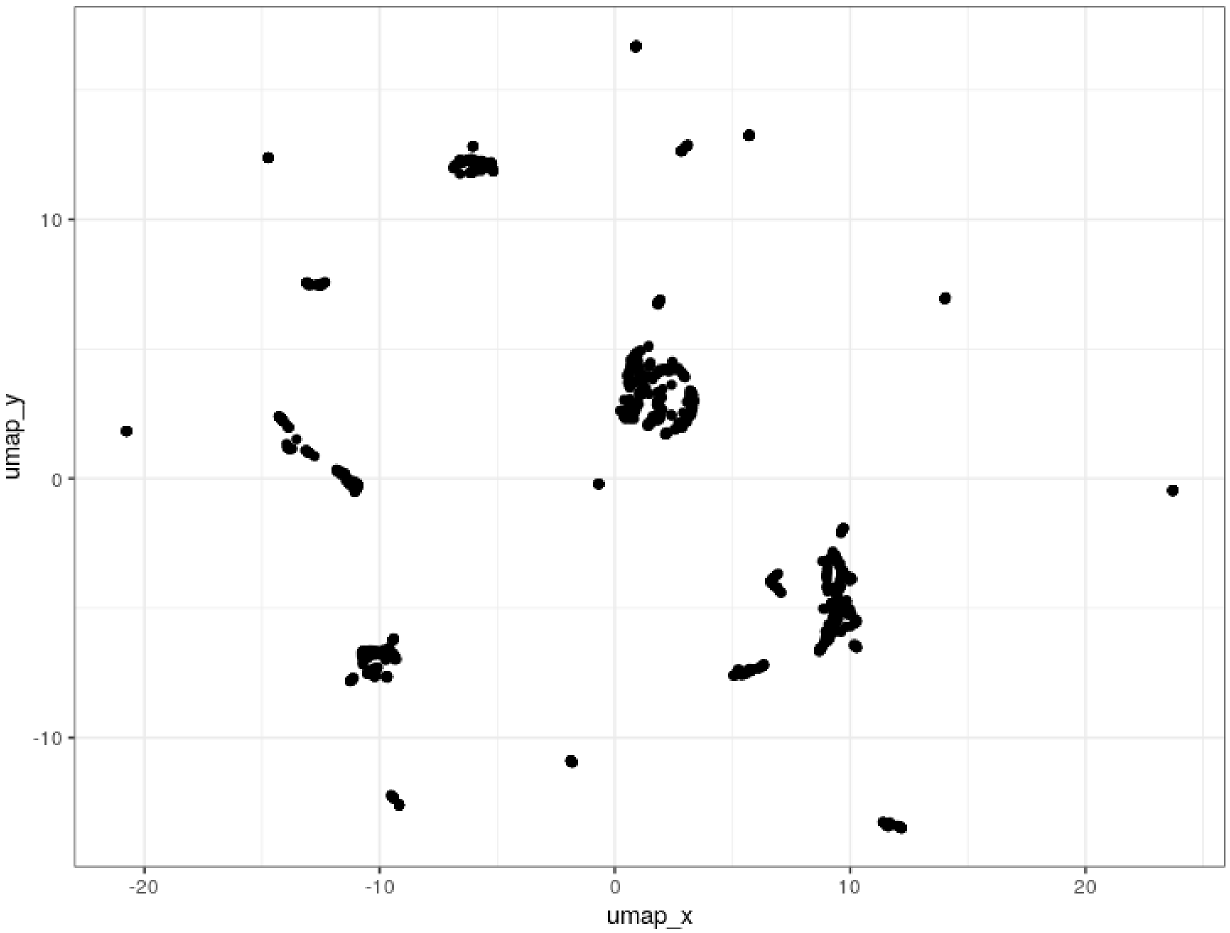

Figure 16: UMAP visualization for MASH data

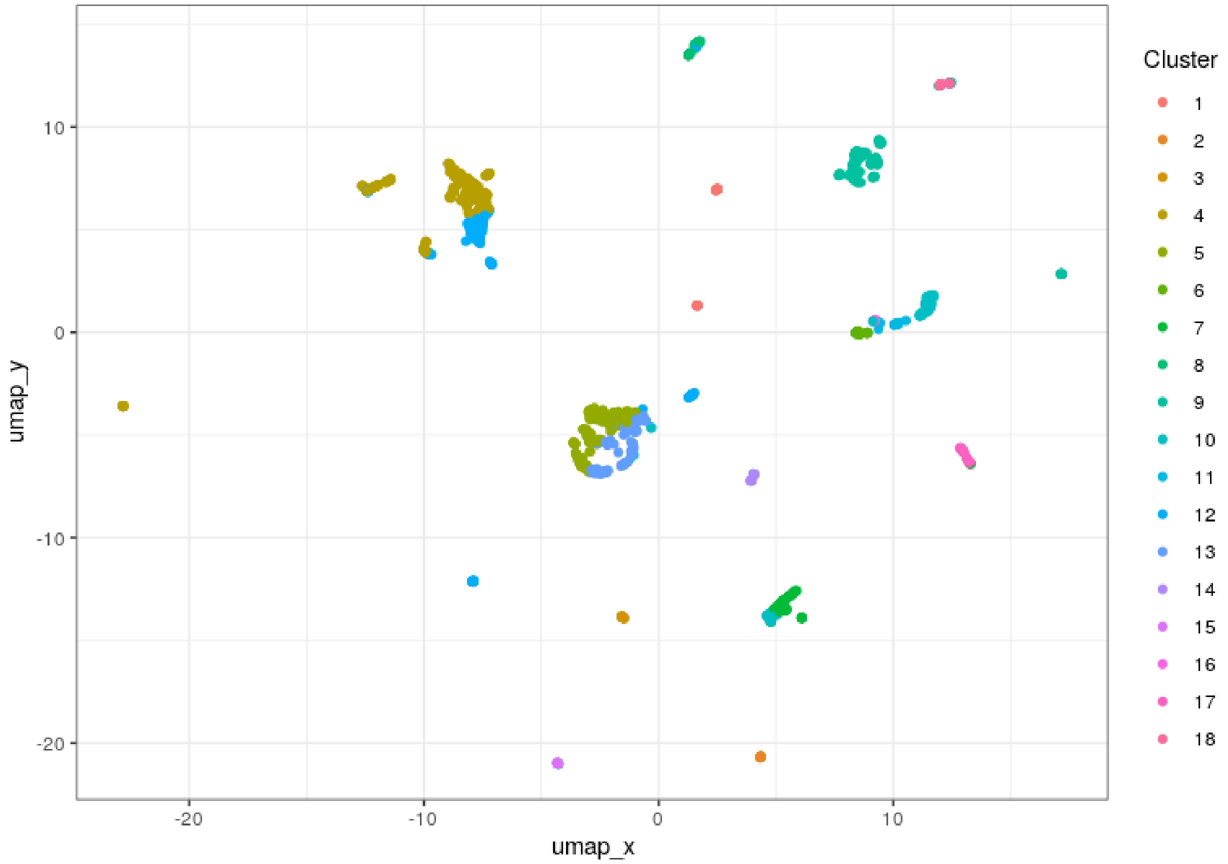

Figure 17: UMAP visualization with cluster definition

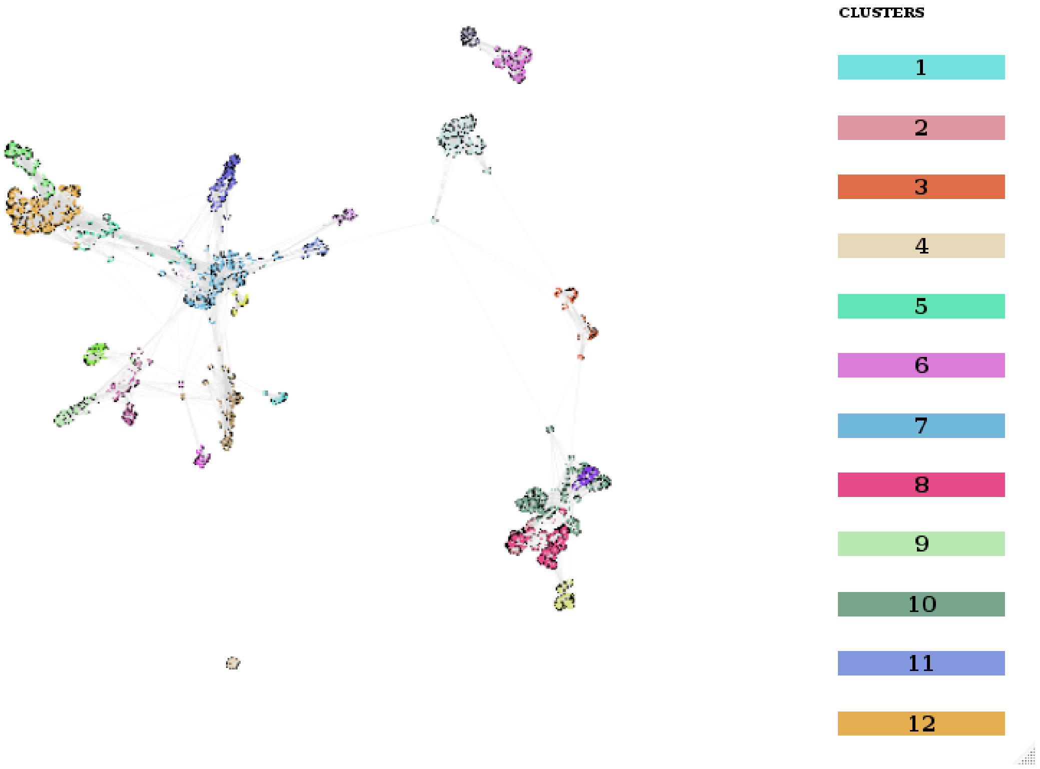

Figure 18: K-NNN with clustering results

```r
  filter(pident > 0.95 ) %>%
  #remove all hits with E-Value greater than 1e-6
  filter(evalue < 1e-6) %>%
  #select only the best hit for each protein-genome.
  group_by(Genome,Protein) %>%
  arrange(desc(bits)) %>%
  slice_head()
```

PATO also includes a function to create a heatmap with the annotation:

```r
heatmap_of_annotation(annotation %>%
                        filter(DataBase =="AbR"), #We select only "AbR" results
                      min_identity = 0.99)
```

Annotation HeatMap

Or to visualize it as a network.

```r
network_of_annotation(annotation %>%
                        filter(DataBase =="AbR"), min_identity = 0.99) %>%
  export_to_gephi("annotation_Network")
```

[https://doi.org/10.6084/m9.figshare.13121795]

#### Outbreak / Transmission / Description

One of the main goals in microbial genomics is to identify outbreaks or transmission events among different sources. Basically, that we want to do, is to find the same strain (or very similar) if different subjects/sources. Commonly that means to find two (or more) strains differing in less than a few SNPs between them.

For this example we are going to use the dataset published in [https://msphere.asm.org/content/5/1/e00704-19] (you can download the data set at https://doi.org/10.6084/m9.figshare.13482435.v1).

In this paper, authors performed whole-genome comparative analyses on 60 *E. coli* isolates from soils and fecal sources (cattle, chickens, and humans) in households of rural Bangladesh. PATO implements different ways to identify Outbreak/Transmission events. The most standard way was to calculate the core-genome, to create the phylogenetic tree and to establish the (average) number of SNPs difference between the strains (or eventually the ANI). We can use four different alternatives:

- MASH similarity
- Core-genome + snp_matrix (roary-like)
- Core_snp_genome + snp_matrix (snippy-like)
- Snps-pairwise (most expensive but most accurate)

First we download the data and decompress in a folder. In my case *~/example PATO* folder. We set that folder as a working directory.

```r
setwd("~/examplePATO/)
```

The *Montealegre et al.* dataset contains 60 genomes in GFF3 format including the sequence in FASTA format. We load the genomes into PATO.

```r
##Creates a file-list
gff_files <- dir("~/examplesPATO/", pattern = "\\.gff", full.names =T)
##Load the genomes
gffs <- load_gff_list(gff_files)
```

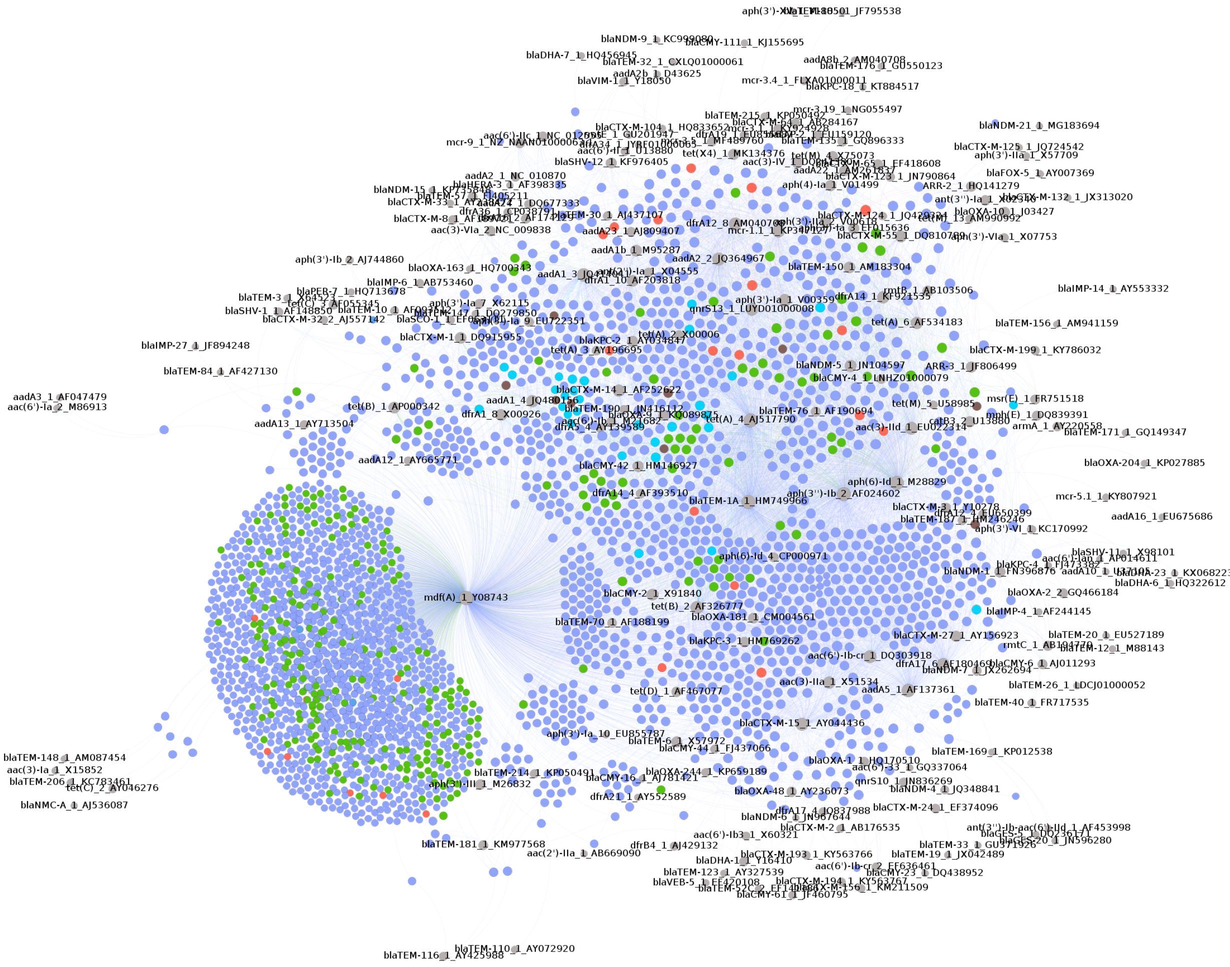

Figure 19: Gephi visualization of the network of AbR genes

We also create a file with the real names of each sample. Using *bash* command line, we are going to extract the names of each sample.

```
grep -m 1 'strain' *.gff
  >| sed 's/;/\t/g' | sed 's/:/\t/g'
  >| cut -f1,22,23 | sed 's/nat-host=.*\t//'
  >| sed 's/strain=//' > names.txt
```

Now we load the names into R.

```
#Load the files
strain_names <- read.table("names.txt")%>%
  #Rename the columns
  rename(Genome = V1, Name = V2) %>%
  #delete the '-' character
  mutate(Name = gsub("-","",Name)) %>%
  #Extract the first 4 character as Sample name
  mutate(Sample = str_sub(Name,1,4)) %>%
  #Extract the final characters as Source
  mutate(Source = str_sub(Name,5)) %>%
  #remove the final part of the filename
  mutate(Genome = str_replace(Genome,"_genomic.gff",""))
```

The samples are designed with the name of the household number and the source (H: Human, C: Cattle, CH: Chicken, S: Soil). For example HH15CH is the householder 15 and the source is a chicken feces.

##### Mash Similarity.

The fastest (and easy way) to find high similarities is using the MASH distance.

```
mash <- mash(gffs, type ="wgs", n_cores = 20)
mash_tree <- similarity_tree(mash,method = "fastme") %>%
  phangorn::midpoint()
```

Now using *ggtree* we can add the names info into the tree.

```
# remove '_genomic.fna' from the tip.labels to fix with the strain_names table
mash_tree$tip.label <- gsub("_genomic.fna","",mash_tree$tip.label)
ggtree(mash_tree) %<+%
  strain_names +
  geom_tippoint(aes(color=Source)) +
  geom_tiplab(aes(label = Sample))
```

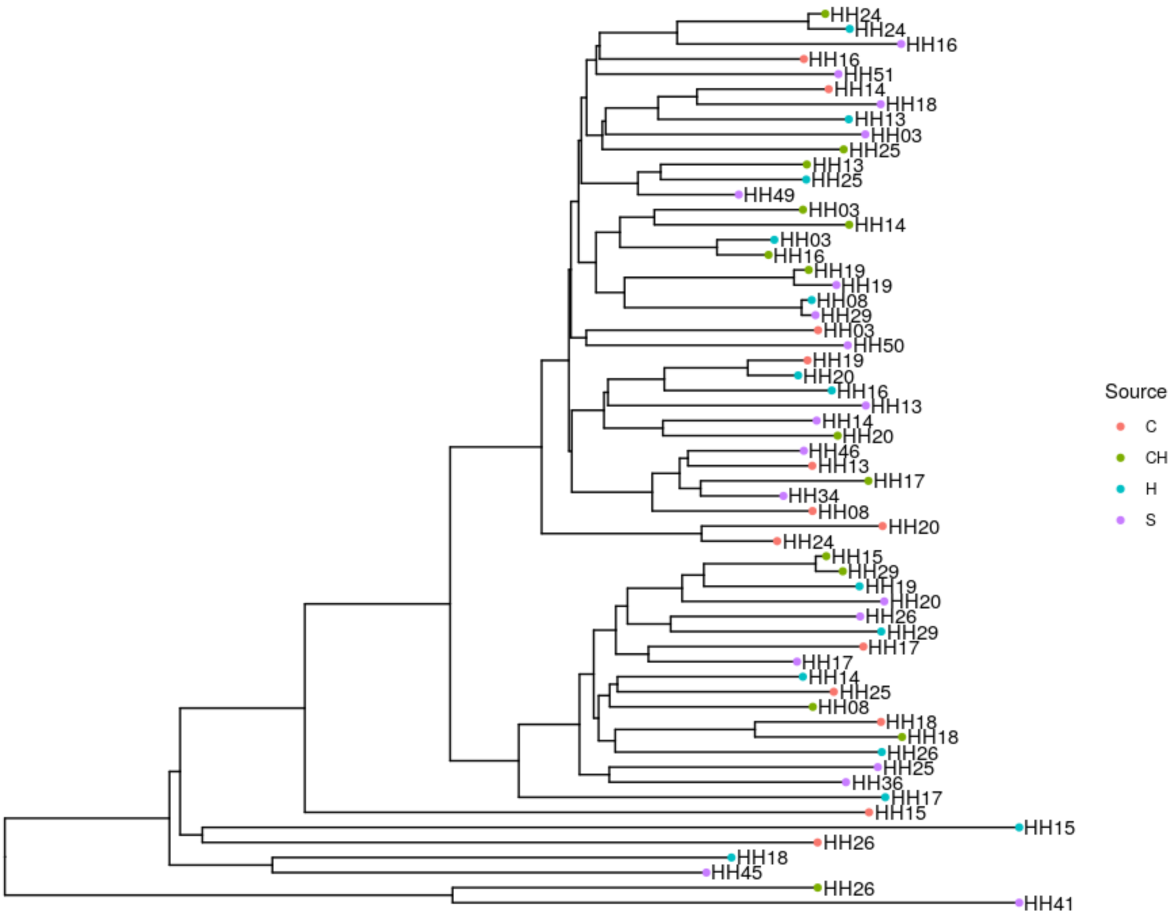

We can inspect the similarity values. Remember that the MASH distance is equivalent to $D_{mash} = 1 - \frac{ANI}{100}$

```
mash$table %>%
  #Remove extra characters
  mutate(Source = gsub("_genomic.fna","",Source)) %>%
  mutate(Target = gsub("_genomic.fna","",Target)) %>%
  #Add real names to Source
  inner_join(strain_names %>%
               rename(Source2 = Name) %>%
               select(Genome,Source2),
             by= c("Source" = "Genome")) %>%
  #Add real names to Target
  inner_join(strain_names %>%
               rename(Target2 = Name) %>%
               select(Genome,Target2),
             by= c("Target" = "Genome")) %>%
  #Remove diagonal
  filter(Source != Target) %>%
  #Compute the ANI
  mutate(ANI = (1-Dist)*100) %>%
  #Filter hits higher than 99.9% identity
  filter(ANI > 99.9) %>%
  #Transform to data.frame to show all digits of ANI
```

```
  as.data.frame()

  Source2 Target2      ANI
1   HH08H   HH29S 99.94674
2  HH15CH  HH29CH 99.91693
3  HH29CH  HH15CH 99.91693
4   HH29S   HH08H 99.94674
```

We must take into account that MASH uses the whole genome to estimate the genome distance.That means that the accessory genomes (plasmids, transposons phages. . . ) are considered to calculate the distance.

Usually we use the number of SNPs as a measure of clonal relatedness . We are going to compare the MASH way to the SNPs approaches.

##### Core-genome + snp_matrix (roary-like)

In this case we are going to use the PATO pipeline to compute the core-genome and the number of SNPs among the samples.

```
mm <- mmseqs(gffs, type = "nucl")
core <- core_genome(mm,type = "nucl", n_cores = 20)
export_core_to_fasta(core,file = "pato_roary_like.fasta")

#Externally compute the phylogenetic tree
system("fasttreeMP -nt -gtr pato_roary_like.fasta > pato_roary_like.tree")

#Import and rooting the tree
pato_roary_tree <- ape::read.tree("pato_roary_like.tree") %>%
  phangorn::midpoint()
#Fix the tip labels
pato_roary_tree$tip.label <- gsub("_genomic.ffn","",pato_roary_tree$tip.label)

#Draw the tree with the annotation.
ggtree(pato_roary_tree) %<+%
  strain_names +
  geom_tippoint(aes(color=Source)) +
  geom_tiplab(aes(label = Sample))
```

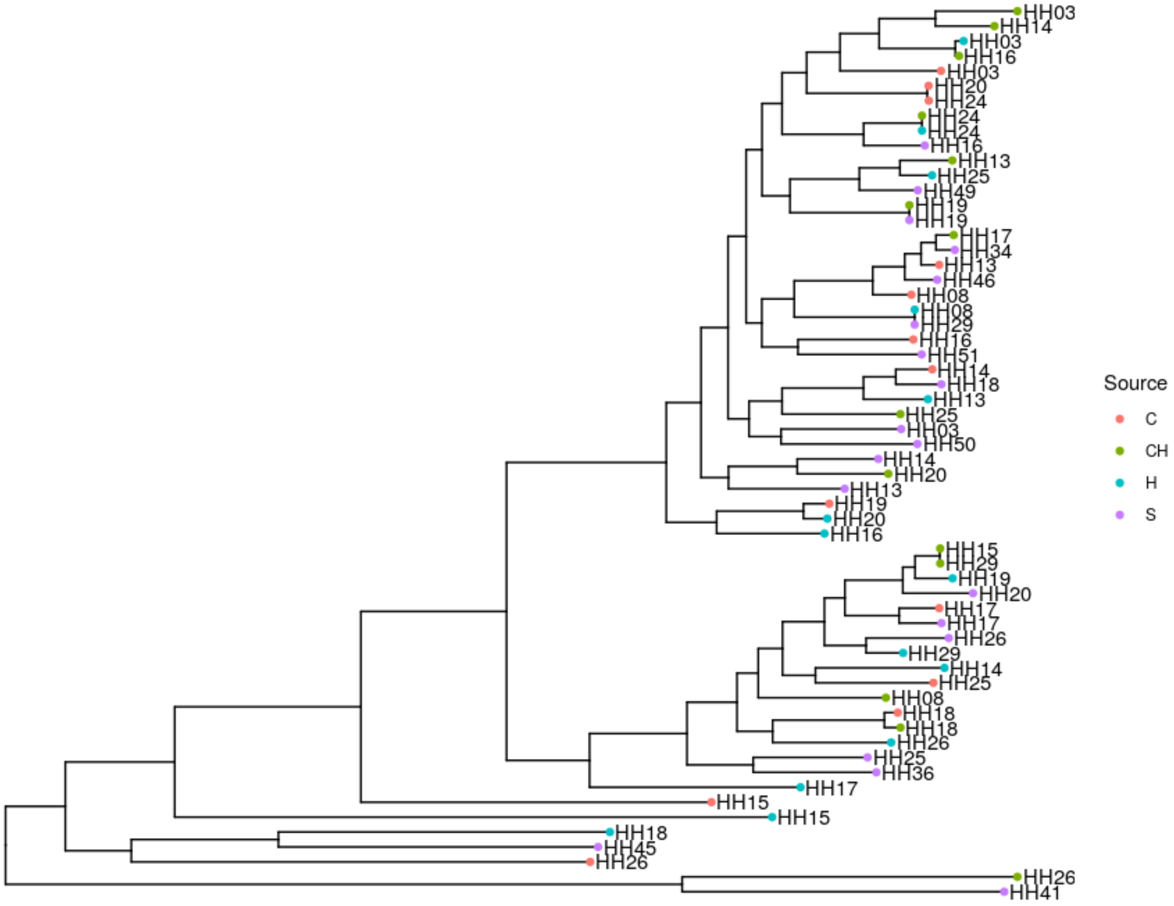

Now we can compute the SNPs matrix

```
#compute the SNP matrix removing columns with gaps
pato_roary_m = core_snps_matrix(core, norm = T, rm.gaps = T)

#Fix the colnames and rownames to put the real names.

colnames(pato_roary_m) <-
  rownames(pato_roary_m) <-
  gsub("_genomic.ffn","",rownames(pato_roary_m))
tmp = pato_roary_m %>%
  as.data.frame() %>%
  rownames_to_column("Genome") %>%
  inner_join(strain_names) %>%
  column_to_rownames("Name") %>%
  select(-Genome,-Source,-Sample) %>%
  as.matrix()
colnames(tmp) = rownames(tmp)

#Plot the matrix as a heatmap
pheatmap::pheatmap(tmp,
         #To Display only values lower than 200 SNPs
         display_numbers = matrix(ifelse(tmp < 200,tmp,""), nrow(tmp)),
```

```
         number_format = '%i',
         annotation_row = strain_names %>%
           select(-Genome,-Sample) %>%
           column_to_rownames("Name"),
         annotation_col = strain_names %>%
           select(-Genome,-Sample) %>%
           column_to_rownames("Name"),
         show_rownames = T,
         show_colnames = T,
)
```

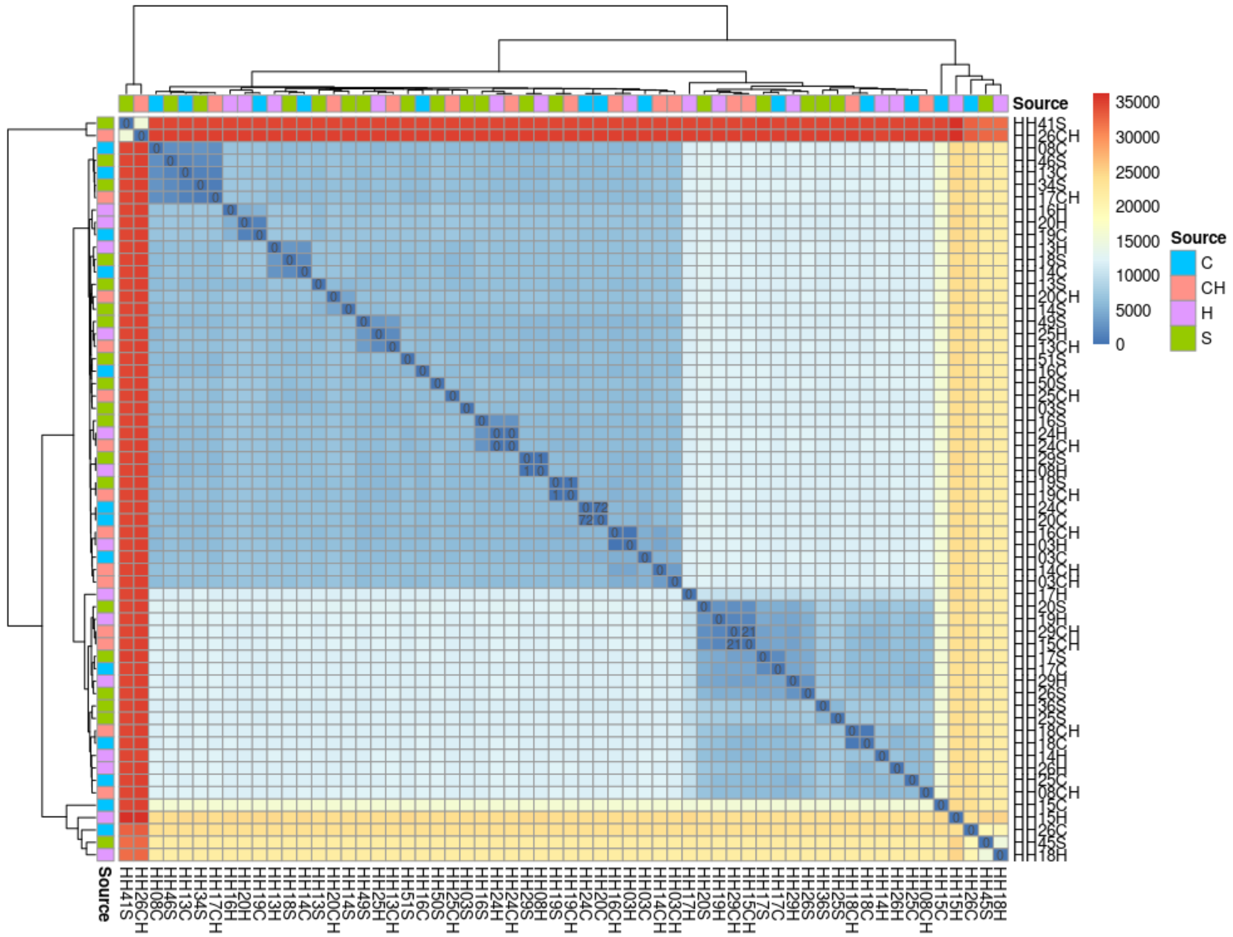

We can manually inspect the matrix in the console.

```
tmp %>%
  as.data.frame() %>%
  rownames_to_column("Source") %>%
  pivot_longer(-Source,names_to = "Target", values_to = "SNPs") %>%
  filter(Source != Target) %>%
  filter(SNPs < 200)

# A tibble: 10 x 3
   Source Target  SNPs
   <chr>  <chr>  <dbl>
 1 HH29S  HH08H      1
```

```
 2 HH29CH HH15CH    21
 3 HH24H  HH24CH     0
 4 HH24CH HH24H      0
 5 HH24C  HH20C     72
 6 HH20C  HH24C     72
 7 HH19S  HH19CH     1
 8 HH19CH HH19S      1
 9 HH15CH HH29CH    21
10 HH08H  HH29S      1
```

We find three possible transmissions HH24 from Human to Chicken (or vice versa), from HH29 Soil to HH08 Human (or vice versa) and Human HH19 to chicken (or vice versa) that have less than 4 SNPs/ Mb . PATO compute the number of SNPs normalized by the length of the alignment in Mb
(i.e. the core-genome)

##### Core_snp_genome + snp_matrix (snippy-like)

Another option to find possible transmission is using the *core_snp_genome()* function. In this case, all the genomes are aligned against a reference. *core_snp_genome()* find the common regions among all the samples and extract the SNPs of those regions. This approach is very similar to Snippy (https://github.com/tseemann/snippy) pipeline.

```
core_s <- core_snp_genome(gffs,type = "wgs")
export_core_to_fasta(core_s,file = "pato_snippy_like.fasta")
#Externally compute the phylogenetic tree
system("fasttreeMP -nt -gtr pato_snippy_like.fasta > pato_snippy_like.tree")

#Import and rooting the tree
pato_snippy_tree <- ape::read.tree("pato_snippy_like.tree") %>%
  phangorn::midpoint()
#Fix the tip labels
pato_snippy_tree$tip.label <- gsub("_genomic.fna","",pato_snippy_tree$tip.label)

#Draw the tree with the annotation.
ggtree(pato_snippy_tree) %<+%
  strain_names +
  geom_tippoint(aes(color=Source)) +
  geom_tiplab(aes(label = Sample))
```

Now the SNPs matrix

```
colnames(pato_snippy_m) <-
  rownames(pato_snippy_m) <-
  gsub("_genomic.fna","",rownames(pato_snippy_m))
tmp = pato_snippy_m %>%
  as.data.frame() %>%
  rownames_to_column("Genome") %>%
  inner_join(strain_names) %>%
  column_to_rownames("Name") %>%
  select(-Genome,-Source,-Sample) %>%
  as.matrix()

colnames(tmp) = rownames(tmp)
```

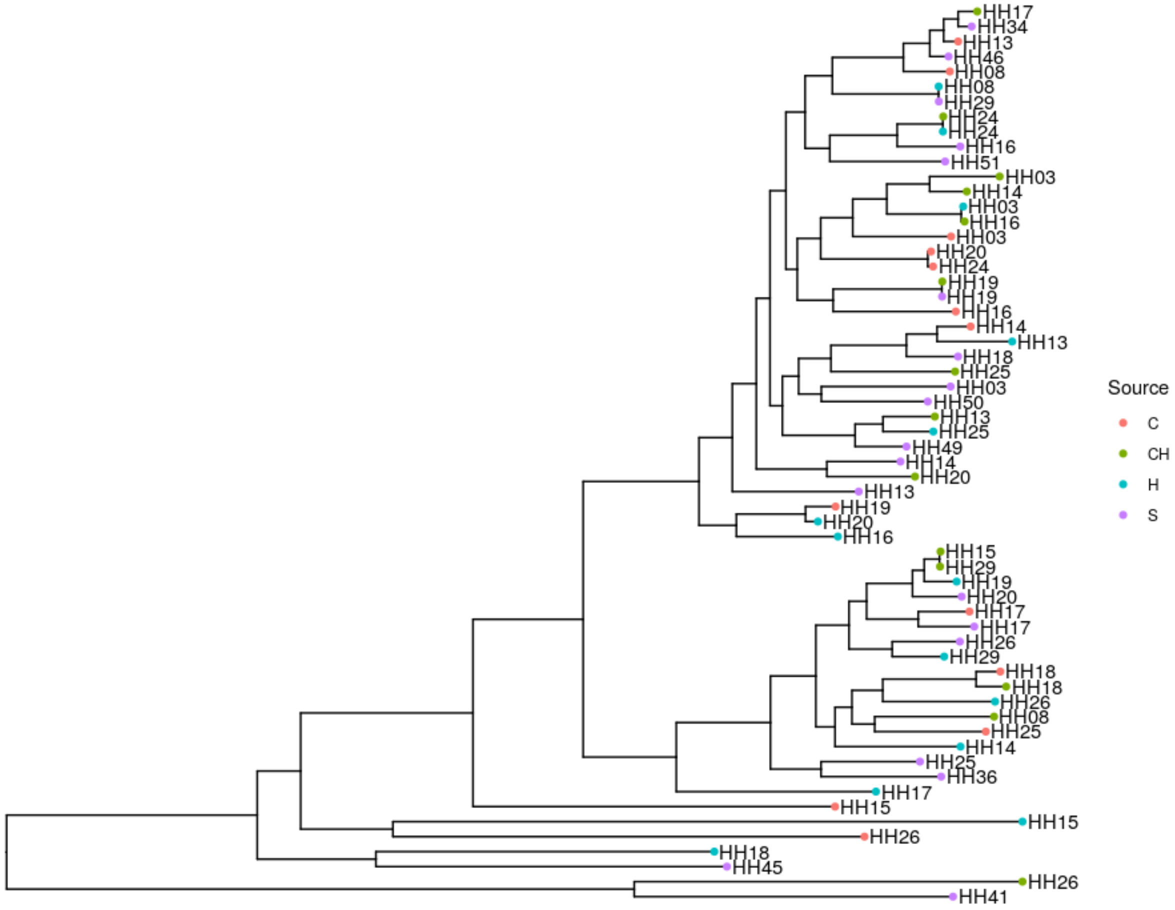

Figure 20: Phylogenetic tree of Core SNP genome

```r
pheatmap(tmp,
         display_numbers = matrix(ifelse(tmp < 200,tmp,""), nrow(tmp)),
         number_format = '%i',
         annotation_row = strain_names %>%
           select(-Genome,-Sample) %>%
           column_to_rownames("Name"),
         annotation_col = strain_names %>%
           select(-Genome,-Sample) %>%
           column_to_rownames("Name"),
         show_rownames = T,
         show_colnames = T,
         )
```

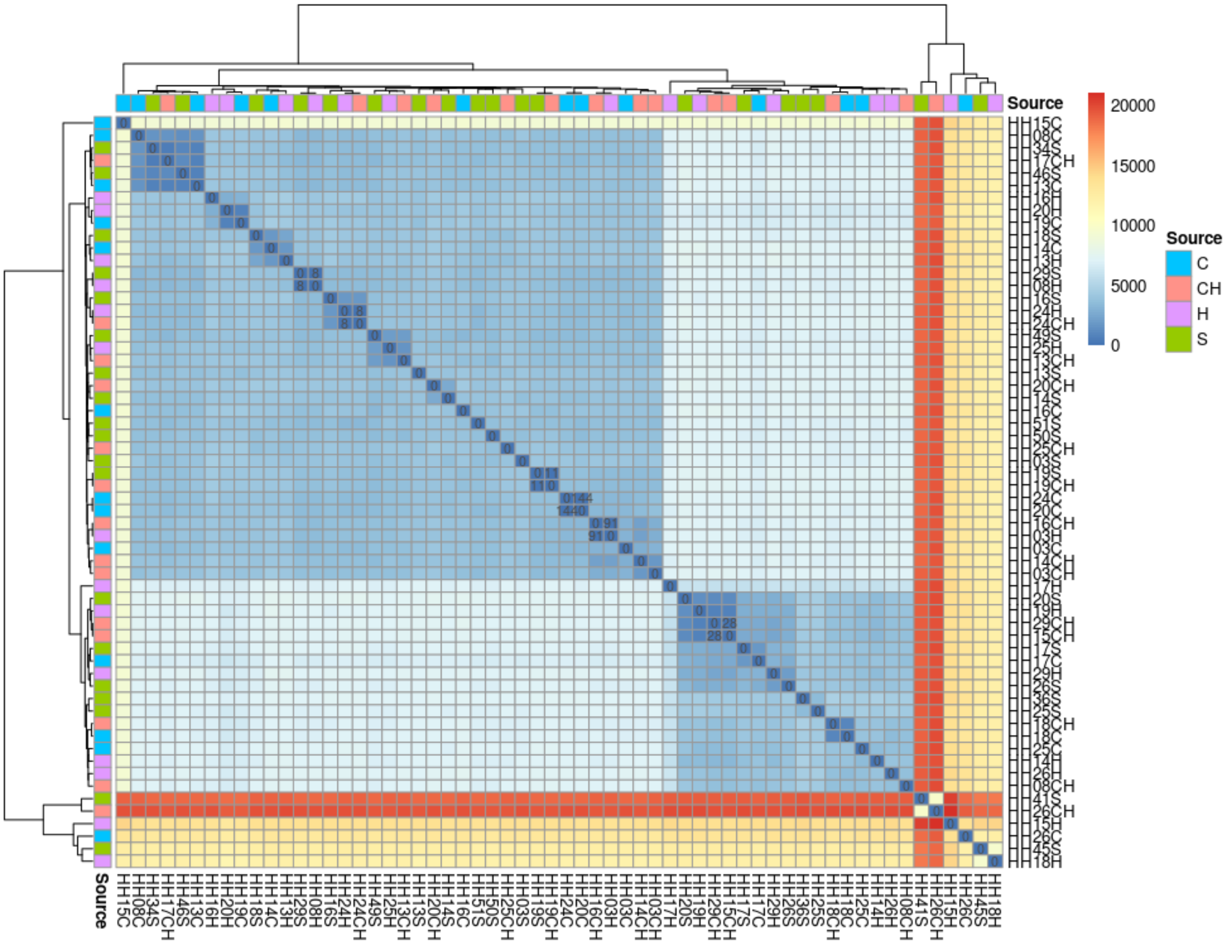

Figure 21: HeatMap of the *core-snp-genome* SNPs matrix

And a manual inspection

```r
tmp %>%
  as.data.frame() %>%
  rownames_to_column("Source") %>%
  pivot_longer(-Source,names_to = "Target", values_to = "SNPs") %>%
  filter(Source != Target) %>%
  filter(SNPs < 200)
```

```
# A tibble: 12 x 3
   Source Target  SNPs
   <chr>  <chr>  <dbl>
 1 HH29S  HH08H      8
 2 HH29CH HH15CH    28
 3 HH24H  HH24CH     8
 4 HH24CH HH24H      8
 5 HH24C  HH20C    144
 6 HH20C  HH24C    144
 7 HH19S  HH19CH    11
 8 HH19CH HH19S     11
 9 HH16CH HH03H     91
10 HH15CH HH29CH    28
11 HH08H  HH29S      8
12 HH03H  HH16CH    91
```

With this approach the number of SNPs increase but the pair are the same HH24H<->HH24CH, HH29S<->HH08H and HH19S<->HH19CH

##### Snps-pairwaise (more computationally expensive but more accurate)

The last two approaches share a limitation: they depend on the dataset structure. In both cases we are counting the number of SNPs of the core-genome, but the core-genome depends on the dataset. If the dataset contains an outlier or if it is heterogeneously diverse, the core-genome could be narrowed and underestimate the number of SNPs, even normalized by the alignment length. PATO implements a pairwise computation of SNPs number with the functions *snps_pairwaise()*. This function can be computationally intense because it performs an alignment among all the sequences $O(n^2)$ in comparison with *core_snp_genome()* that performs $O(n)$. We do not recommend using a dataset larger than 100 genomes (that means 10.000 alignments).

```
pw_normalized <- snps_pairwaise(gffs, type = "wgs",norm = T, n_cores = 20)

tmp = pw_normalized %>%
  as.data.frame() %>%
  rownames_to_column("Genome") %>%
  inner_join(strain_names) %>%
  column_to_rownames("Name") %>%
  select(-Genome,-Source,-Sample) %>%
  as.matrix()
colnames(tmp) = rownames(tmp)

pheatmap(tmp,
         display_numbers = matrix(ifelse(tmp < 200,tmp,""), nrow(tmp)),
         number_format = '%i',
         annotation_row = strain_names %>%
           select(-Genome,-Sample) %>%
           column_to_rownames("Name"),
         annotation_col = strain_names %>%
           select(-Genome,-Sample) %>%
           column_to_rownames("Name"),
         show_rownames = T,
         show_colnames = T,
)
```

And the manual inspection reveals that

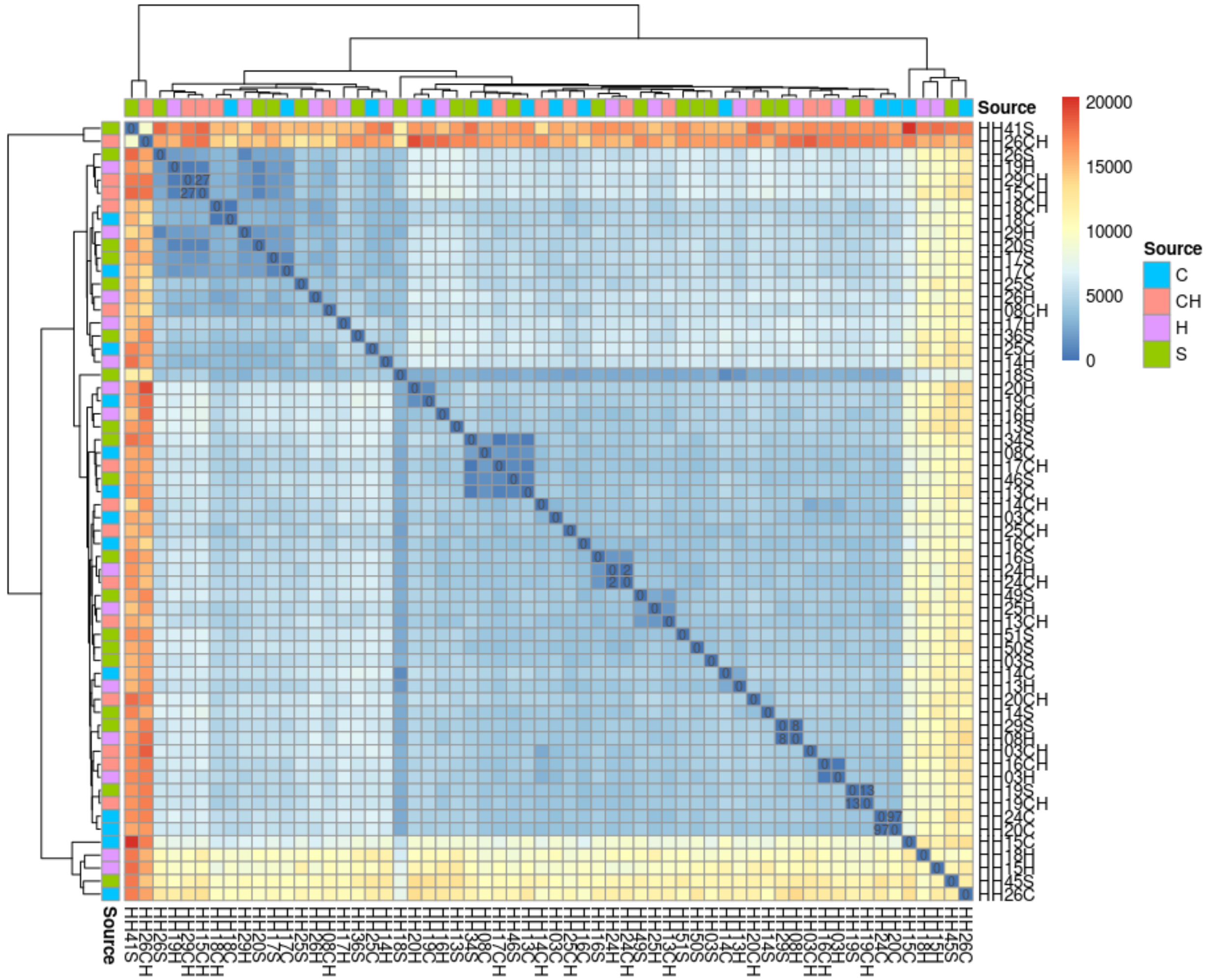

Figure 22: HeatMap of the pairwaise SNP Matrix

```r
tmp %>%
  as.data.frame() %>%
  rownames_to_column("Source") %>%
  pivot_longer(-Source,names_to = "Target", values_to = "SNPs") %>%
  filter(Source != Target) %>%
  filter(SNPs < 200)

# A tibble: 10 x 3
   Source Target  SNPs
   <chr>  <chr>  <dbl>
 1 HH29S  HH08H      8
 2 HH29CH HH15CH    27
 3 HH24H  HH24CH     2
 4 HH24CH HH24H      2
 5 HH24C  HH20C     97
 6 HH20C  HH24C     97
 7 HH19S  HH19CH    13
 8 HH19CH HH19S     13
 9 HH15CH HH29CH    27
10 HH08H  HH29S      8
```

And that is the real number of SNPs per-megabase, excluding indels.

We can conclude that HH24 and XXXX reflect a recent transmission event while HH29<->HH08 and HH19 appears as a more distant transmission.

these results are in disagreement with the MASH results , p probably due to "dispensable" or "accesory" elements (plasmids, phages, ICEs etc...) which are not taken into account in any of the approaches based on SNPs differences .

#### Pangenomes Analysis

PATO is also designed to analyze the relationship among pangenomes of different (or not) species. The goal is to analyze cluster of genomes (pangenomes) among them as individual elements. In this example, we analyze 49.591 genomes of the bacterial phylum Firmicutes available at the NCBI database. For that kind of large datasets, it is recommended to use the specific pipeline for pangenomes which first clusters the genomes into pangenomes ( e.g. cluster of species, intra-species phylogroups or sequence types (ST)). The diversity of each cluster depends on the *distance*parameter. The pipeline creates homogeneous (in phylogenetic distance) clusters. Then, the pipeline produces an `accnet` object so all the above described function can be used with this object as a common `accnet` object. The pipeline also includes parameters to set the minimum amount of genomes to be considered as a pangenome, and the minimum frequency of a protein/gene family to be included in a pangenome.

```r
res <- pangenomes_from_files(files,distance = 0.03,min_pange_size = 10,
                             min_prot_freq = 2)

export_to_gephi(res,"/storage/tryPATO/firmicutes/pangenomes_gephi")
```

In this case, we produce a gephi table to visualize the accnet network of the dataset. To annotate the network we use the NCBI assembly table to give a species name to each pangenome cluster.

```r
assembly <- data.table::fread("ftp://ftp.ncbi.nlm.nih.gov/genomes/refseq/bacteria/assembly_summary.txt"
                             sep = "\t",
                             skip = 1,
                             quote = "")
```

```r
colnames(assembly) = gsub("# ","",colnames(assembly))

annot = res$members %>%
  #mutate(file = basename(path)) %>%
  separate(path,c("GCF","acc","acc2"),sep = "_", remove = FALSE) %>%
  unite(assembly_accession,GCF,acc,sep = "_") %>%
  left_join(assembly) %>%
  separate(organism_name,c("genus","specie"), sep = " ") %>%
  group_by(pangenome,genus,specie) %>%
  summarise(N = n()) %>%
  distinct() %>% ungroup() %>%
  group_by(pangenome) %>%
  slice_head() %>%
  mutate(ID = paste("pangenome_",pangenome,"_rep_seq.fasta",
                    sep = "",collapse = ""))

annot <- annot %>%
  mutate(genus = gsub("\\[","",genus)) %>%
  mutate(genus = gsub("\\]","",genus)) %>%
  mutate(specie = gsub("\\[","",specie)) %>%
  mutate(specie = gsub("\\]","",specie)) %>%
  unite("Label",genus,specie, remove = F)

write_delim(annot,"../pangenomes_gephi_extra_annot.tsv",delim = "\t",
            col_names = TRUE)
```

We determined the species name as the most frequent species for each cluster. Then using Gephi we visualize the network.

Another way to visualize the results is a heatmap of the sharedness among the pangenoms. PATO includes the function sharedness to visualize this result. To visualize the `data.table` result from `sharedness()` function we can use the package `pheatmap`. As the input value of `pheatmap` can only be a matrix, first, we must transform our result into a matrix:

```r
sh <-  sharedness(res)
sh  <-  sh %>% as.data.frame() %>%
  rownames_to_column("ID") %>%
  inner_join(annot) %>%
  unite(Name,genus,specie,pangenome, sep = "_") %>%
  unite("Row_n",Label,N) %>%
  select(-ID)%>%
  column_to_rownames("Row_n")

colnames(sh) = rownames(sh)
```

Now we can execute `pheatmap()` with our pangenome dataset.

```r
pheatmap::pheatmap(log2(sh+1),
                   clustering_method = "ward.D2",
                   clustering_distance_cols = "correlation",
                   clustering_distance_rows = "correlation")
```

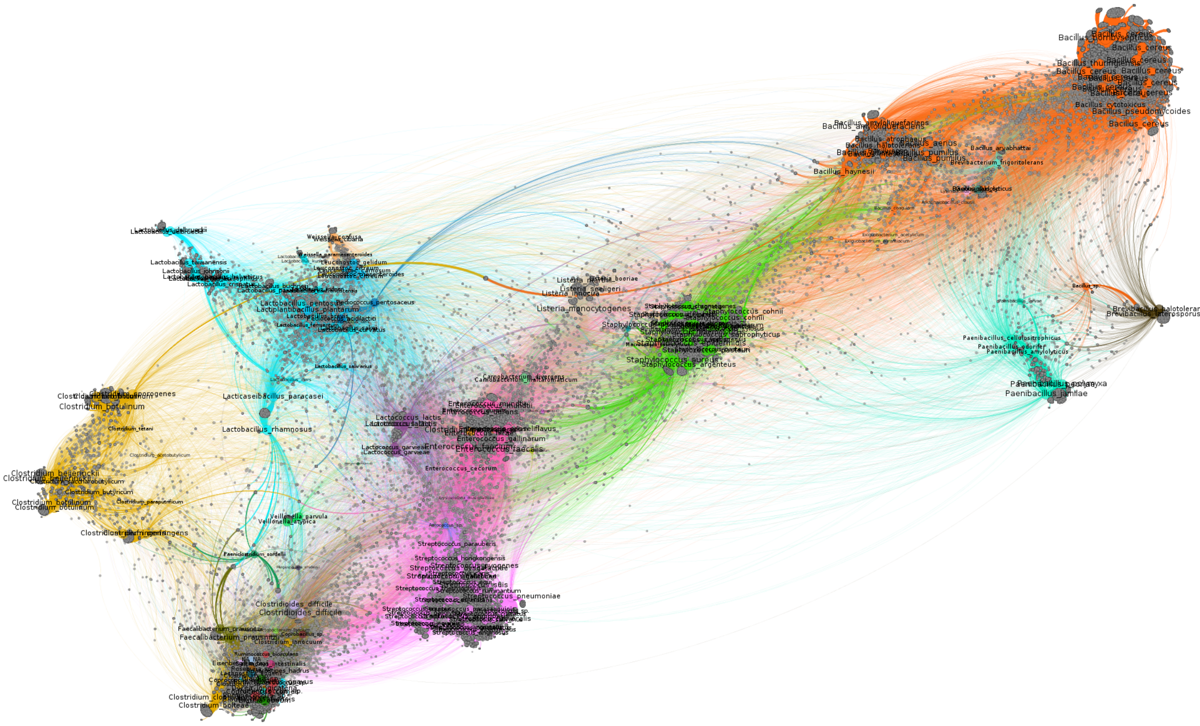

Figure 23: Pangenomes network of firmicutes: Gephi visualization

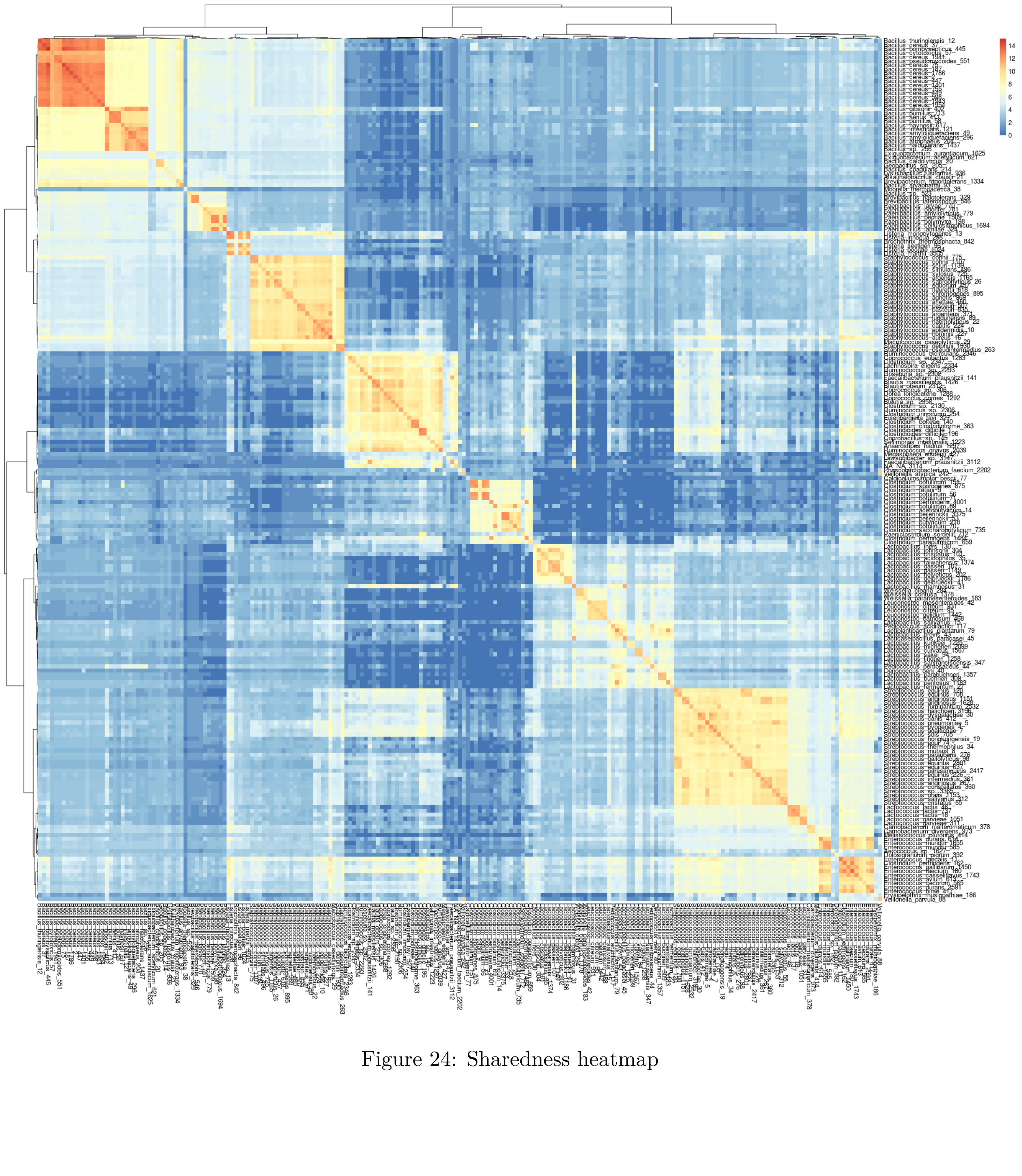

Figure 24: Sharedness heatmap

#### Benchmarking

Although there is not software or package that can be fully compared with PATO we have compared PATO with two of the most used software Roary and Snippy. Both can be used to compute a core genome alignment. Roary computes a gene-by-gene core-genome alignment and Snippy uses a mapping approach to build a core snp genome.

We have compared the performance of both packages against PATO using the montealegre et al. dataset.

To measure the performance we used *microbenchmark* package. We have tried to reproduce the same steps for each pipeline as PATO performs these actions using different functions.

```
library(microbenchmark)

bench <- microbenchmark(

  "roary_time" = {system("roary -p 24 -o roary_ex -e --mafft *.gff")},

  "pato_like_roary_time"= {
    gffs <- load_gff_list(gff_files)
    mm <- mmseqs(gffs, type = "nucl")
    acc <- accnet(mm)
    core <- core_genome(mm,type = "nucl", n_cores = 24)
    core_p <- core_plots(mm,steps = 10,reps = 5)
    export_accnet_aln(file = "accesory_aln.fa",acc)
    system("fasttreeMP -nt accesory_aln.fa > accessory_aln.tree")
  },

  "pato_like_snippy_time" = {
    unlink(gffs$path,recursive = T)
    gffs <- load_gff_list(gff_files)
    core_s <- core_snp_genome(gffs,type = "wgs",
            ref = "~/examplesPATO/fna/GCF_009820385.1_ASM982038v1_genomic.fna")
    export_core_to_fasta(core_s,file = "pato_snippy_like.fasta")
  },

  "snippy_time" = {
    setwd("~/examplesPATO/fna")

    # That is my $PATH. Do not try to execute.
    #To execute Snippy from R you need to define the $PATH
    Sys.setenv(PATH ="/usr/local/bin:/usr/bin:/bin: ....")

    system("ls *.fna > tmp2")
    system("ls $PWD/*.fna > tmp")
    system("sed -i 's/_genomic.fna//' tmp")
    system("paste tmp tmp2 > list_snippy.txt")
    system("snippy-multi list_snippy.txt --ref
           GCF_009820385.1_ASM982038v1_genomic.fna --cpus 24 > snp.sh")
    system("sh ./snp.sh")
  },
  times = 1
)
```

The benchmarking results was

Unit: *seconds*

| expr | min | lq | mean | median | uq | max | neval |
|---|---|---|---|---|---|---|---|
| roary_time | 6459.1894 | 6459.1894 | 6459.1894 | 6459.1894 | 6459.1894 | 6459.1894 | 1 |
| pato_like_roary_time | 185.4633 | 185.4633 | 185.4633 | 185.4633 | 185.4633 | 185.4633 | 1 |
| pato_like_snippy_time | 149.9739 | 149.9739 | 149.9739 | 149.9739 | 149.9739 | 149.9739 | 1 |
| snippy_time | 3425.9932 | 3425.9932 | 3425.9932 | 3425.9932 | 3425.9932 | 3425.9932 | 1 |

Therefore, PATO (equivalent pipeline) is almost 35x faster than Roary and 23x faster than Snippy for a dataset of 60 genomes in a Dual Intel Xeon with 24 effective cores (12 real cores) and 48 Gb RAM.

If we compare the results of core alignments, Roary creates 2.649.299 bp alignment (including indels) and PATO 2.902.678 bp. If we compare against Snippy then, Snippy creates a 4.680.136 bp alignment and PATO 4.392.982 bp.

Moreover, Roary finds 227.383 SNPs and PATO 253.654. Those SNPs are computed excluding indels. Snippy finds 260.909 and PATO 155.300. We think that difference can be caused by in *core_snps_genome()* functions as they do not count the multi-variant positions (changes of two or more nucleotides in a row). We are working to fix it in future versions of PATO.

If we compare the phylogenetic tree obtained with the same software (Fasttree) we see that

```
library(ape)
library(TreeDist)
library(phangorn)

pato_snippy_tree = read.tree("pato_snippy_like.tree") %>% midpoint()
pato_snippy_tree$tip.label = gsub("_genomic.fna","",pato_snippy_tree$tip.label)

pato_roary_tree = read.tree("pato_roary_like.tree") %>% midpoint()
pato_roary_tree$tip.label = gsub("_genomic.ffn","",pato_roary_tree$tip.label)

snippy_tree = read.tree("snippy.tree") %>% midpoint()
snippy_tree = TreeTools::DropTip(snippy_tree,"Reference")

roary_tree = read.tree("roary.tree") %>% midpoint()
roary_tree$tip.label = gsub("_genomic","",roary_tree$tip.label)

rand_tree = rtree(60, tip.label = snippy_tree$tip.label) %>% midpoint()

mash <- mash(gffs, type ="wgs", n_cores = 20)
mash_tree <- similarity_tree(mash,method = "fastme") %>% midpoint()

mash_tree$tip.label = gsub("_genomic.fna","",mash_tree$tip.label)

trees <- c(random_tree = rand_tree,
           snippy_tree = snippy_tree,
           roary_tree = roary_tree,
           pato_core = pato_roary_tree,
           pato_core_snp = pato_snippy_tree,
           mash = mash_tree)
```

```r
CompareAll(trees,RobinsonFoulds)%>%
  as.matrix() %>%
  pheatmap::pheatmap(.,display_numbers = T)
```

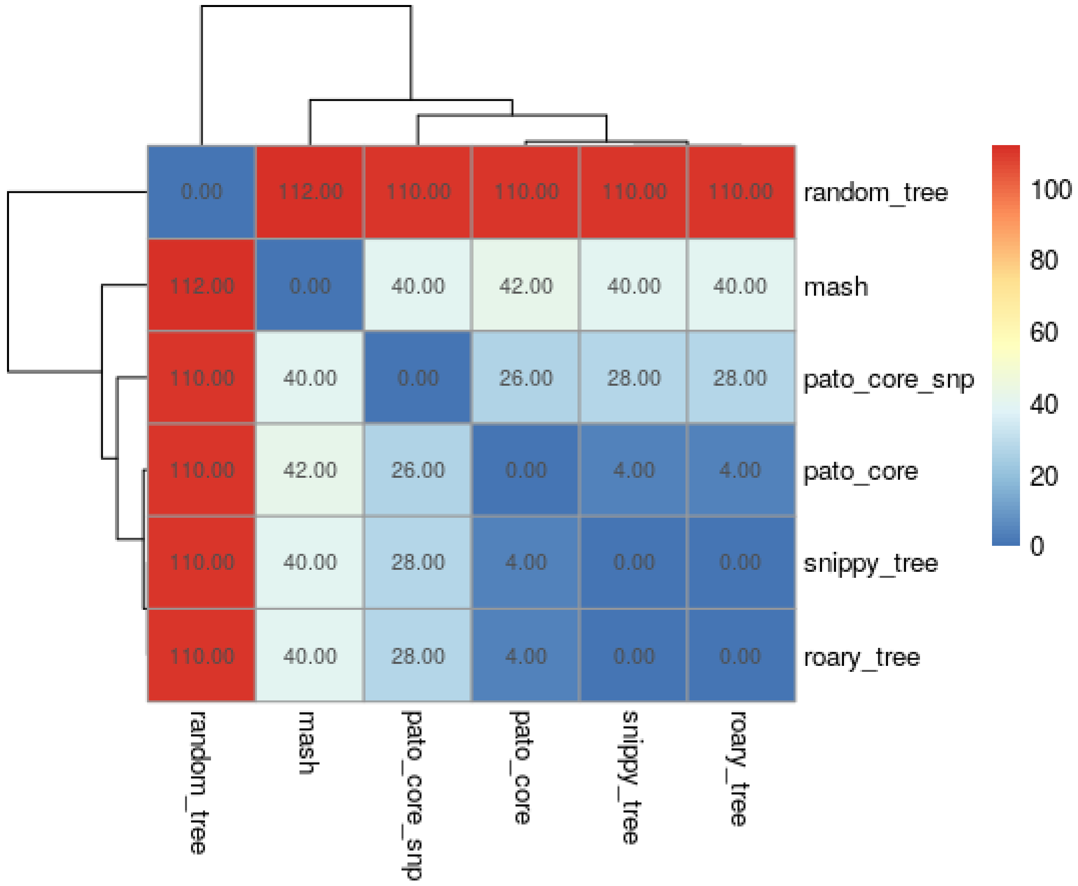

Figure 25: HeatMap of the Robinson Foulds distances

We compare the trees using *TreeDist* package. This package implements Robison-Foulds method to measure phylogenetic tree similarities.

While Roary and Snippy obtain the same phylogenetic tree, PATO roary-like obtain a very similar tree and PATO snippy-like obtain an accurate tree but not exactly the same. In comparison to the MASH tree, the alignment of all these methods are quite similar among them. We also include a random tree as a reference to an unrelated tree.

#### Session Info

```
> sessionInfo()
R version 3.6.3 (2020-02-29)
Platform: x86_64-pc-linux-gnu (64-bit)
Running under: Debian GNU/Linux 10 (buster)

Matrix products: default
BLAS:   /usr/lib/x86_64-linux-gnu/blas/libblas.so.3.8.0
```

```
LAPACK: /usr/lib/x86_64-linux-gnu/lapack/liblapack.so.3.8.0

locale:
 [1] LC_CTYPE=es_ES.UTF-8       LC_NUMERIC=C               LC_TIME=es_ES.UTF-8
LC_COLLATE=es_ES.UTF-8     LC_MONETARY=es_ES.UTF-8
 [6] LC_MESSAGES=es_ES.UTF-8    LC_PAPER=es_ES.UTF-8       LC_NAME=C
LC_ADDRESS=C               LC_TELEPHONE=C
[11] LC_MEASUREMENT=es_ES.UTF-8 LC_IDENTIFICATION=C

attached base packages:
[1] stats     graphics  grDevices utils     datasets  methods   base

other attached packages:
 [1] pheatmap_1.0.12      stringdist_0.9.6.3   microseq_2.1.2       rlang_0.4.8
data.table_1.13.2    TreeDist_1.2.1
 [7] microbenchmark_1.4-7 phangorn_2.5.5       ape_5.4-1            ggtree_2.0.4
pato_1.0.1           forcats_0.5.0
[13] stringr_1.4.0        dplyr_1.0.2          purrr_0.3.4          readr_1.3.1
tidyr_1.1.2          tibble_3.0.4
[19] ggplot2_3.3.2        tidyverse_1.3.0

loaded via a namespace (and not attached):
  [1] Rtsne_0.15            colorspace_1.4-1      ellipsis_0.3.1
mclust_5.4.6          XVector_0.26.0        base64enc_0.1-3
  [7] fs_1.5.0              rstudioapi_0.11       farver_2.0.3
bit64_4.0.5           fansi_0.4.1           lubridate_1.7.9
 [13] xml2_1.3.2            codetools_0.2-16      R.methodsS3_1.8.1
doParallel_1.0.16     jsonlite_1.7.1        broom_0.7.0
 [19] cluster_2.1.0         dbplyr_1.4.4          R.oo_1.24.0
uwot_0.1.8            shiny_1.5.0           BiocManager_1.30.10
 [25] compiler_3.6.3        httr_1.4.2            rvcheck_0.1.8
backports_1.1.10      lazyeval_0.2.2        assertthat_0.2.1
 [31] Matrix_1.2-18         fastmap_1.0.1         cli_2.1.0
later_1.1.0.1         htmltools_0.5.0       tools_3.6.3
 [37] igraph_1.2.6          gtable_0.3.0          glue_1.4.2
V8_3.2.0              fastmatch_1.1-0       Rcpp_1.0.5
 [43] cellranger_1.1.0      vctrs_0.3.4           Biostrings_2.54.0
nlme_3.1-144          crosstalk_1.1.0.1     iterators_1.0.13
 [49] gbRd_0.4-11           rbibutils_2.0         openxlsx_4.2.2
rvest_0.3.6           mime_0.9              miniUI_0.1.1.1
 [55] lifecycle_0.2.0       zlibbioc_1.32.0       scales_1.1.1
hms_0.5.3             promises_1.1.1        parallel_3.6.3
 [61] RColorBrewer_1.1-2    curl_4.3              memoise_1.1.0
dtplyr_1.0.1          stringi_1.5.3         randomcoloR_1.1.0.1
 [67] S4Vectors_0.24.4      foreach_1.5.1         tidytree_0.3.3
BiocGenerics_0.32.0   zip_2.1.1             manipulateWidget_0.10.1
 [73] Rdpack_2.1            pkgconfig_2.0.3       parallelDist_0.2.4
lattice_0.20-40       labeling_0.4.2        treeio_1.10.0
 [79] htmlwidgets_1.5.2     bit_4.0.4             tidyselect_1.1.0
magrittr_1.5          R6_2.4.1              IRanges_2.20.2
 [85] generics_0.0.2        DBI_1.1.0             pillar_1.4.6
haven_2.3.1           withr_2.3.0           modelr_0.1.8
 [91] crayon_1.3.4          utf8_1.1.4            grid_3.6.3
```

```
readxl_1.3.1            blob_1.2.1              threejs_0.3.3
 [97] reprex_0.3.0            digest_0.6.26           webshot_0.5.2
xtable_1.8-4            R.cache_0.14.0          httpuv_1.5.4
[103] dbscan_1.1-5            R.utils_2.10.1          TreeTools_1.4.1
openssl_1.4.3           RcppParallel_5.0.2      stats4_3.6.3
[109] munsell_0.5.0           askpass_1.1             quadprog_1.5-8
```
